# Perilipin 3 promotes the formation of membrane domains enriched in diacylglycerol and lipid droplet biogenesis proteins

**DOI:** 10.1101/2022.09.14.507979

**Authors:** Rasha Khaddaj, Roger Schneiter

## Abstract

Lipid droplets (LDs) serve as intracellular stores of energy-rich neutral lipids. LDs form at discrete sites in the endoplasmic reticulum (ER) and they remain closely associated with the ER during lipogenic growth and lipolytic consumption. Their hydrophobic neutral lipid core is covered by a monolayer of phospholipids, which harbors a specific set of proteins. This LD surface is coated and stabilized by perilipins, a family of soluble proteins that specifically target LDs from the cytosol. We have previously used chimeric fusion proteins between perilipins and integral ER membrane proteins to test whether proteins that are anchored to the ER bilayer could be dragged onto the LD monolayer. Expression of these chimeric proteins induces repositioning of the ER membrane around LDs. Here, we test the properties of membrane-anchored perilipins in cells that lack LDs. Unexpectedly, membrane-anchored perilipins induce the formation of crescent-shaped ER membrane domains that have LD-like properties. They are stained by lipophilic dyes and harbor LD marker proteins. In addition, these ER domains are enriched in diacylglycerol (DAG) and in ER proteins that are important for early steps of LD biogenesis, including seipin and Pex30. These ER membrane domains dissolve upon induction of neutral lipid synthesis and the formation of nascent LDs. Formation of these membrane domains *in vivo* requires DAG, and we show that perilipin 3 (PLIN3) binds to liposomes containing DAG *in vitro*. Taken together, these observations indicate that perilipin not only serve to stabilize the surface of mature LDs but that they are likely to exert a more active role in early steps of LD biogenesis at ER subdomains enriched in DAG, seipin, and neutral lipid biosynthetic enzymes.

## Introduction

Most cells use neutral lipids as an efficient anhydrous store of metabolic energy. Neutral lipids, mainly triacylglycerol (TAG) and steryl esters (STE) are concentrated and shielded from the aqueous environment by being packed into intracellular lipid droplets (LDs). The lipidic hydrophobic core of these droplets is enclosed by an unusual phospholipid monolayer, and functionalized by a set of proteins, which specifically localize to the LD surface. While LDs mainly serve as a store of energy and provide a pool of lipid precursors for rapid membrane expansion, they also serve to store proteins, function as sites of virus assembly, and buffer an endoplasmic reticulum (ER) stress response. LD formation and turnover are thus closely integrated within the overall energy metabolism of an organism and the homeostatic control of the lipid composition of membrane-delineated compartments. Consequently, LD function is associated with prevalent pathologies including obesity, atherosclerosis, insulin resistance and fatty liver disease (Krahmer et al., 2013; Welte & Gould, 2017; Gluchowski et al., 2017; Xu et al., 2018; Olzmann & Carvalho, 2018).

LDs are formed from the ER membrane, where the neutral lipid biosynthetic enzymes, such as the main TAG biosynthetic enzymes, diacylglycerol acyltransferases (DGAT1 and DGAT2 in humans, Dga1 and Lro1 in yeast), and the sterol acyltransferases (ACATs in humans, Are1 and Are2 in yeast), reside (Czabany et al., 2007; Turkish & Sturley, 2009). The neutral lipids produced by these enzymes are released into the ER bilayer, where they have low solubility and coalesce to form neutral lipid lenses/blisters (Hamilton et al., 1983; Khandelia et al., 2010). These lipid lenses then grow in size, and emerge towards the cytoplasm as nascent LDs, which further grow and mature in their protein and lipid composition (Thiam & Forêt, 2016; Thiam & Ikonen, 2021).

More recent results indicate that LD biogenesis in the ER membrane is both temporally and spatially highly regulated and occurs at specialized ER subdomains, which contain the LD biogenesis protein seipin, the seipin associated proteins LDAF1/promethin in mammals and Ldo16/45 in yeast, the membrane-shaping protein Pex30, the fat storage-inducing transmembrane (FIT) proteins, the ER tether Mdm1, and a protein phosphatase complex composed of Nem1 and Spo7 (Kassan et al., 2013; Hariri et al., 2019; Cao et al., 2019; Choudhary et al., 2020; Henne et al., 2020; Renne et al., 2020; Schneiter & Choudhary, 2022). LDs are thus formed from ER subdomains and remain functionally and structurally closely associated with the ER to exchange both lipids and proteins (Zehmer et al., 2009; Jacquier et al., 2011; Wilfling et al., 2013; Choudhary et al., 2015; Cottier & Schneiter, 2022).

Proteins can target the surface of LDs either from the cytoplasmic space or through the ER membrane (Olarte et al., 2021). Perilipins (PLINs) form a family of soluble proteins that specifically associate with the LD surface (Greenberg et al., 1991). Mammalian PLINs and their yeast ortholog Pet10, are cytosolic in the absence of LDs, but are recruited onto the LD surface upon induction of their biogenesis (Brasaemle, 2007; Bulankina et al., 2009; Jacquier et al., 2013; Gao et al., 2017). PLINs are abundant proteins that are thought to act as a scaffold to stabilize the LD surface, and to regulate lipolytic activation of lipases (Kimmel & Sztalryd, 2016; Sztalryd & Brasaemle, 2017; Itabe et al., 2017).

Most PLIN family members are composed of three domains, an N-terminal PAT domain, followed by a repeat of amphipathic helices (11-mer repeats) and a C-terminal domain that adopts a four 4-helix bundle conformation. The amphipathic repeat segments that are present in PLIN family members are similar to those found in apolipoproteins and the Parkinson’s disease-associated alpha-synuclein. They are important for targeting the LD surface possibly by recognizing lipid packing defects in the LD monolayer (Bussell & Eliezer, 2003; Rowe et al., 2016; Copic et al., 2018; Giménez-Andrés et al., 2018; Ajjaji et al., 2019; Giménez-Andrés et al., 2021). The 4-helix bundle, on the other hand, has structural similarity to apolipoprotein E, and fragments liposomes into disc-like structures *in vitro* (Hickenbottom et al., 2004; Bulankina et al., 2009). PLINs may have more active function in LD biogenesis as they may bind and possibly sequester lipids, particularly diacylglycerol (DAG), already in the ER and thereby promote LD biogenesis (Skinner et al., 2009; Jacquier et al., 2013).

We have recently developed membrane-anchored LD probes by fusing PLIN3 to the cytosolic or the ER luminal ends of ER residential integral membrane proteins (Khaddaj et al., 2022). Cytosolic exposure of PLIN3 in these fusion proteins induced tight wrapping of the ER membrane around LDs, as monitored by a split-GFP readout. These results indicate that membrane spanning ER proteins were excluded from reaching the LD surface, suggesting that a barrier between the ER bilayer and the LD monolayer limits the access of integral membrane proteins to the LD surface (Khaddaj et al., 2022).

Here, we analyze the properties of membrane-anchored PLIN3 fusion proteins in cells lacking LDs. We show that membrane-proximal PLINs induce the formation of crescent-shaped membrane domains in the ER. Formation of these domains is lipid-dependent and they colocalize with an ER-DAG sensor. Remarkably, PLIN-induced membrane domains have the properties of LDs as they can be labelled with neutral-lipid specific fluorescent dyes and they colocalize with LD marker proteins. They also recruit ER proteins that promote LD formation such as seipin or Pex30, indicating that PLINs have membrane-organizing properties and might concentrate or stabilize neutral lipids within a phospholipid membrane. Consistent with this proposition, PLIN3 binds DAG *in vitro* and specifically associates with liposomes containing DAG.

## Results

### Membrane-anchored perilipin 3 induces the formation of crescent-shaped ER domains in cells lacking LDs

We have previously reported that the expression of chimeric fusion proteins between membrane-spanning integral ER membrane proteins such as Sec61, a multispanning transmembrane protein and subunit of the ER translocon (Deshaies & Schekman, 1987), or Wbp1, a single spanning membrane protein and component of the oligosaccharyl transferase complex required for N-linked glycosylation (te Heesen et al., 1992), and the mammalian LD-targeted protein perilipin 3 (PLIN3/TIP47; Wolins et al., 2001; Bulankina et al., 2009) results in wrapping of the ER membrane around LDs (Khaddaj et al., 2022) (Fig. 1A). In the absence of PLIN3, Wbp1-GFP and Sec61-GFP display homogenous ER localization and no significant colocalization with the LD marker Erg6-mCherry (Khaddaj et al., 2022) (Fig. 1B). When fused to PLIN3, however, Wbp1-GFP-PLIN3 and Sec61-GFP-PLIN3 colocalize with Erg6-mCherry at the rim of LDs in cells supplemented with oleic acid to increase the size and number of LDs (Khaddaj et al., 2022) (Fig. 1C). Tight wrapping of the ER membrane around LDs in cells expressing the membrane-anchored PLIN3 creates the impression of a *bona fide* LD-localization of the fusion proteins. However, split-GFP readouts have indicated that these membrane-anchored PLIN3 reporter proteins induce a rearrangement of the ER membrane around LDs, rather than localize on the LD monolayer (Khaddaj et al., 2022) (Fig. 1A).

**Figure 1.**
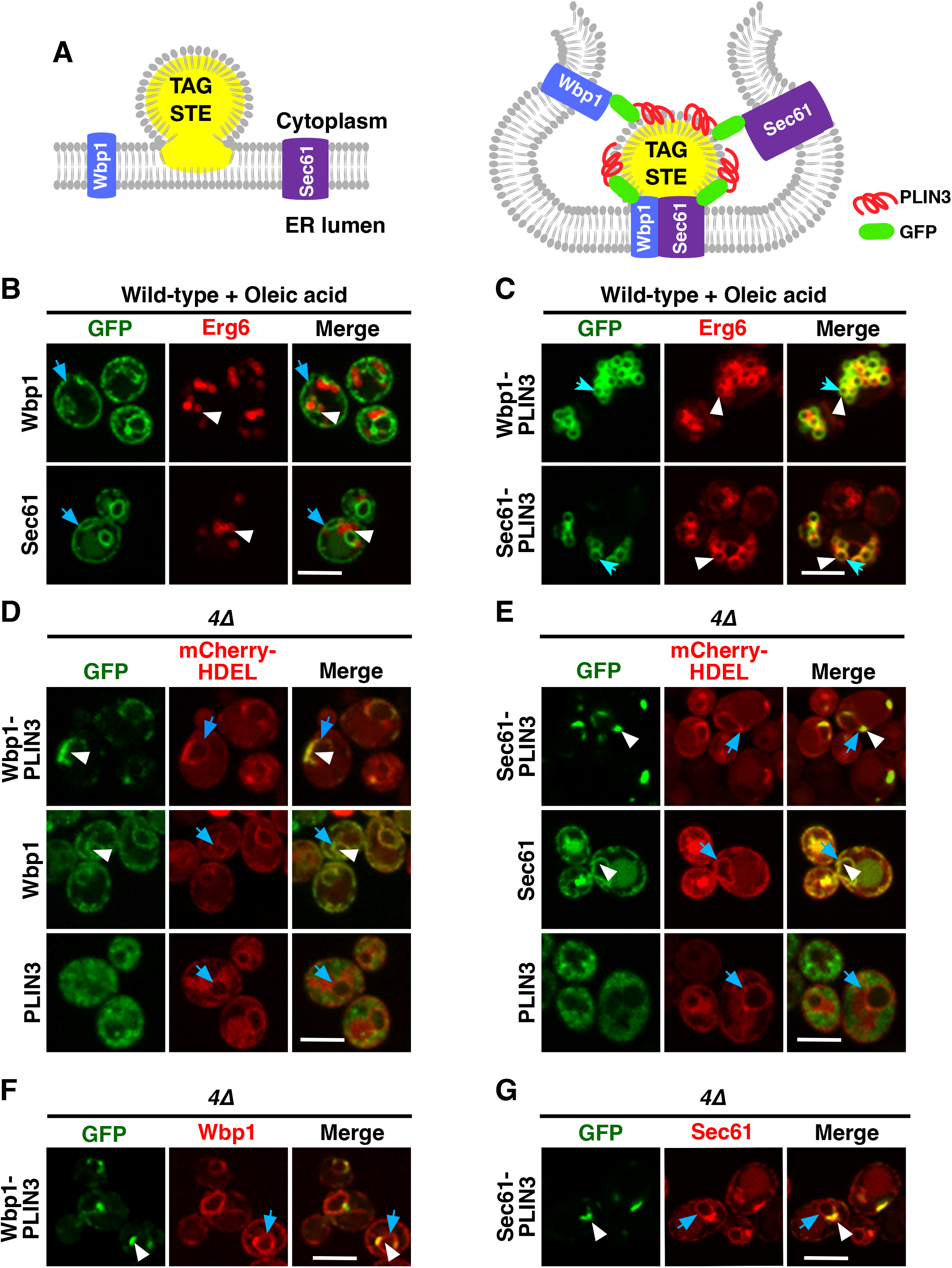
Membrane-anchored PLIN3 induce formation of ER domains in cells lacking LDs. A) Schematic drawing of the structure and localization of the membrane-anchored reporter proteins used in this study. The ER residential integral membrane proteins Wbp1, a single spanner and subunit of the glycosyltransferase, or Sec61, a polytopic transmembrane protein and subunit of the translocon, were fused to a GFP reporter (green oval) and the LD-targeting perilipin 3 (PLIN3, red helix). Native Sec61 and Wbp1, i.e., when not fused to PLIN3, are localized to the ER membrane in cells containing LDs (left-hand panel). LDs are shown as globular structure filed with triacylglycerol, TAG, and steryl esters, STE, indicated in yellow. When appended to PLIN3, however, Wbp1-GFP-PLIN3 and Sec61-GFP-PLIN3, induce wrapping of the ER around LDs and thus appear to localize to the rim of LDs (right-hand panel). B) ER localization of Wbp1-GFP and Sec61-GFP (not fused to PLIN3) in wild-type cells. Cells coexpressing GFP-tagged Wbp1 or Sec61 together with Erg6-mCherry were grown in medium containing oleic acid and analyzed by confocal microscopy. Blue arrows point to the peripheral ER, white arrowheads mark LDs. Scale bar, 5 μm. C) LD-localization of the membrane-anchored PLIN3. Wild-type cells coexpressing Wbp1-GFP-PLIN3 or Sec61-GFP-PLIN3 with Erg6-mCherry were grown in medium containing oleic acid and analyzed by confocal microscopy. White arrowheads and blue arrows indicate colocalization of the markers at the rim of LDs in the merged image. Scale bar, 5 μm. D, E) In cells lacking LDs, membrane-anchored PLIN3 localize to crescent-shaped ER domains. In quadruple mutant cells lacking LDs (*4*Δ, *are1*Δ *are2*Δ *dga1*Δ *lro1*Δ) Wbp1-GFP-PLIN3 (D) and Sec61-GFP-PLIN3 (E) localize to crescent-shaped and punctate domains in the ER (white arrowheads) and colocalize with the ER luminal marker mCherry-HDEL (blue arrows). Wbp1-GFP and Sec61-GFP, lacking PLIN3, display homogenous ER localization (white arrowheads) and colocalize with mCherry-HDEL (blue arrows). Soluble PLIN3-GFP is localized to the cytosol in cells lacking LDs, no significant labelling of the ER membrane (blue arrows) is observed when PLIN3 is not anchored to an integral membrane protein. Scale bar, 5 μm. F, G) Colocalization of native and PLIN3-containing versions of Wbp1 and Sec61. The native mScarlet-tagged versions of Wbp1 (F) and Sec61 (G) (blue arrows) display some enrichment in membrane domains containing Wbp1-GFP-PLIN3 or Sec61-GFP-PLIN3 (white arrowheads), respectively. Scale bar, 5 μm.

In cells lacking LDs due to the deletion of the neutral lipid biosynthetic genes: *DGA1* and *LRO1* for the synthesis of TAG and *ARE1* and *ARE2* for the synthesis of STE (*4*Δ mutant cells), otherwise LD-localized proteins such as the sterol biosynthetic enzyme Erg6 or the TAG lipase Tgl3 are localized in the ER membrane (Jacquier et al., 2011; Schmidt et al., 2013). To examine whether such a homogenous ER localization would also be observed with the membrane-anchored PLIN3, we transformed *4*Δ mutant cells with plasmids expressing Wbp1-GFP-PLIN3 and Sec61-GFP-PLIN3. In these *4*Δ mutant cells, Wbp1-GFP-PLIN3 and Sec61-GFP-PLIN3 labelled crescent-like domains within the ER membrane, as shown by partial colocalization with the ER luminal marker mCherry-HDEL (Fig. 1D, E). Formation of these ER crescents required the presence of PLIN3, because Wbp1-GFP and Sec61-GFP displayed homogenous ER localization and did not induce formation of crescent-shaped ER domains. PLIN3 alone, on the other hand, was cytosolic in these LD-deficient cells, as observed before (Jacquier et al., 2013) (Fig. 1D, E). Formation of crescent-shaped ER domains by membrane-proximal PLIN3 impacted the distribution of other ER proteins to different degrees. While Wbp1-mScarlet was still homogenously distributed over the entire nuclear ER, Sec61-mScarlet displayed some accumulation within the crescent-shaped domains formed by Sec61-GFP-PLIN3 (Fig. 1F, G). Taken together, these results indicate that a membrane-proximal positioning of PLIN3 results in the formation of ER membrane domains in cells that have no LDs.

### ER domains generated by membrane-anchored PLIN3 transform upon induction of LD biogenesis

Given that expression of membrane-anchored PLIN3 results in the formation of crescent-shaped ER domains, we wondered whether these domains were stable or whether they would disperse upon induction of LD formation. To address this, we expressed Wbp1-GFP-PLIN3 and Sec61-GFP-PLIN3 in cells in which TAG synthesis can be induced by a switch of carbon source from glucose to galactose (*GAL-LRO1 are1*Δ *are2*Δ *dga1*Δ). We have previously shown that induction of Lro1 in this strain results in TAG synthesis and the formation of LDs within 30 min of shifting cells to galactose containing medium (Jacquier et al., 2011). Prior to induction of Lro1, the membrane proximal PLIN3 strongly colocalized with the LD marker Erg6-mCherry at ER-crescents (Fig. 2A, B, 0 h). Upon induction of Lro1 expression and hence TAG synthesis, both the membrane-anchored PLIN3 and Erg6 transformed their crescent-shaped localization to label more punctate, LD-like structures (Fig. 2A, B; 1 h time point). After overnight expression of Lro1, Wbp1-GFP-PLIN3 and Sec61-GFP-PLIN3 colocalized with Erg6-mCherry on punctate, circular structures, that are similar to those observed in wild-type cells grown in the presence of oleic acid, suggesting that these marker proteins now colocalize on the rim of LDs (Fig. 2A, B; overnight (ON) time point; Fig1C). This transformation of ER crescents to punctate structures was accompanied by a reduction in the size of the structures labelled by both marker proteins, the membrane-anchored PLIN3 and Erg6 (Fig. 2C). These data indicate that the ER membrane domains formed by membrane-proximal PLIN3 in cells lacking LDs are not caused by irreversible protein aggregation but that these domains transform into LD-like structures upon induction of neutral lipid biosynthesis.

**Figure 2.**
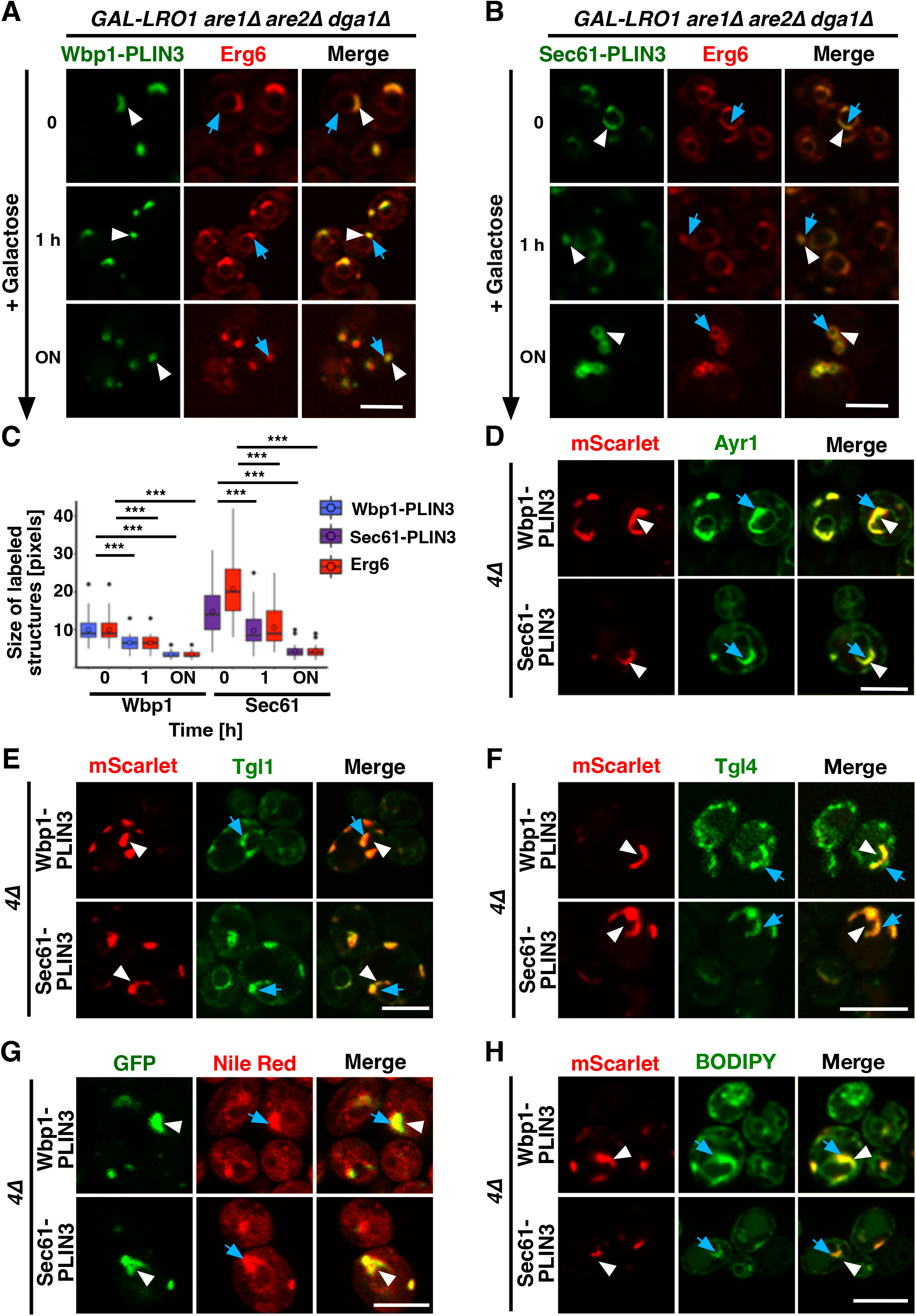
ER domains formed by membrane-anchored PLIN3 transform upon induction of LD biogenesis. A, B) Cells harboring the TAG synthase Lro1 under the control of a galactose inducible promoter (*GAL-LRO1 are1*Δ *are2*Δ *dga1*Δ) and co-expressing Wbp1-GFP-PLIN3 (A), or Sec61-GFP-PLIN3 (B), with Erg6-mCherry were cultivated in glucose medium. At time 0 cells were switched to galactose-containing media to induce expression of Lro1, and hence TAG synthesis, and LD biogenesis. The subcellular distribution of the marker proteins was recorded by confocal microscopy after the indicated periods of time following galactose induction of Lro1 (0, switch to galactose; 1 h, 1 h in galactose; ON, overnight in galactose). The localization of the GFP-tagged membrane-proximal PLIN3 is indicated by white arrowheads, that of Erg6-mCherry by blue arrows. Scale bar, 5 μm. C) Quantification of the size of structures labelled by membrane-proximal PLIN3 or Erg6 during induction of LD biogenesis. The size of the structures labelled by Wbp1-GFP-PLIN3, or Sec61-GFP-PLIN3, compared to Erg6-mCherry at different time points of induction of Lro1 are plotted (N>50 cells). The median is indicated in a box that represents the 25–75th percentile range. The whiskers denote the largest and smallest values with 1.5× of the interquartile range from the hinges of the box. Outliers are depicted by black circles. The significance was assessed using a Wilcoxon signed-rank test with continuity correction. *\*\*\*P*<0.001. D-F) Colocalization between membrane-proximal PLIN3 and LD markers proteins such as Ayr1 (D), Tgl1 (E), or Tgl4 (F). *4*Δ mutant cells expressing Wbp1-mScarlet-PLIN3, or Sec61-mScarlet-PLIN3 together with GFP-tagged versions of the indicated LD markers were imaged by confocal microscopy. Localization of the membrane-proximal PLIN3 is indicated by white arrowheads, that of the LD markers by blue arrows. Scale bar, 5 μm. G, H) Membrane domains induced by the expression of the membrane-proximal PLIN3 are recognized by LD-specific fluorescent dyes Nile Red (G) and BODIPY (H). *4*Δ mutant cells (*are1*Δ *are2*Δ *dga1*Δ *lro1*Δ) expressing GFP- or mScarlet-tagged versions of Wbp1-PLIN3, or Sec61-PLIN3 were incubated with the neutral lipid specific dyes Nile Red or BODIPY and imaged by confocal microscopy. Localization of the membrane-proximal PLIN3 is indicated by white arrowheads, that of the LD-specific fluorescent dyes by blue arrows. Scale bar, 5 μm.

### ER domains formed by the membrane-anchored PLIN3 have LD-like properties

Considering that the ER domains formed by the membrane proximal PLIN3 colocalized with Erg6-mCherry at the perinuclear ER, the site of LD formation, we wondered whether other LD marker proteins would also colocalize within these domains. This is indeed the case as markers of mature LDs such as Ayr1, which encodes a 1-acyldihydroxyacetone-phosphate reductase (Athenstaedt & Daum, 2000), the steryl ester hydrolase Tgl1 (Köffel et al., 2005), or the TAG lipase Tgl4 (Athenstaedt & Daum, 2005), all colocalized with the ER domain induced by the membrane-proximal PLIN3 (Fig. 2D-F).

Remarkably, these crescent-shaped ER membrane domains are not only recognized by proteins as resembling LDs, but they were also stained by commonly used neutral lipid dyes that specifically label LDs such as Nile Red or BODIPY (493/503) (Greenspan et al., 1985; Gocze & Freeman, 1994) (Fig. 2G, H). Because both LD proteins and the neutral lipid dyes selectively stained the PLIN3-containing ER domains, these areas appear to have LD-like properties. Importantly, such membrane domains were not only induced by expression of the membrane-proximal PLIN3 but also when another perilipin family member, PLIN1, was fused to either Wbp1-GFP or Sec61-GFP (Fig. S1A). Thus, the formation of such domains appears to be a general property of membrane-proximal PLINs and their characterization might reveal more general properties of the mode of action of perilipins.

### Formation of ER domains by PLIN3 depends on diacylglycerol

Given that the ER domains, which are induced by expression of membrane-anchored PLIN are recognized both by LD-localized proteins and neutral lipid dyes, we wondered whether these membrane domains are also enriched in specific lipids, in particular DAG. DAG is a neutral lipid that serves as a precursor to TAG formation and that is also needed for the proper biogenesis of LDs (Adeyo et al., 2011; Choudhary et al., 2018). Moreover, upon induction of TAG formation, DAG becomes enriched at specific ER subdomains, where LD biogenesis factors such as seipin promote *de novo* formation of LDs (Choudhary et al., 2020).

To probe for the presence of DAG within the PLIN3 domains, we performed colocalization experiments with a GFP-tagged ER-DAG sensor. This sensor is based on tandem C1 domains from human Protein Kinase D (C1a/b-PKD) fused to GFP, which in turn is fused to the transmembrane region of Ubc6, a tail-anchored ER protein (Fig. 3A). This ER-DAG sensor was previously used to visualize enrichment of DAG at ER subdomains during early steps of LD formation (Choudhary et al., 2018; Choudhary et al., 2020). When expressed in *4*Δ cells that contain versions of the membrane-anchored PLIN3 reporters whose expression was repressed by the addition of doxycycline (Tet^off^), the ER-DAG sensor stained the entire perimeter of the nuclear ER, indicating homogenous distribution of DAG throughout the perinuclear ER (Fig. 3B, C; 0 h time point). Upon induction of Wbp1-mScarlet-PLIN3 or Sec61-mScarlet-PLIN3, however, the ER-DAG sensor strongly colocalized with the crescent-shaped PLIN3 domains (Fig. 3B, C; overnight (ON) time point). A similar colocalization of the ER-DAG sensor with membrane domains formed by expression of membrane-anchored PLIN1 was observed, indicating that the enrichment of DAG within these membrane domains is not specific for PLIN3, but that it appears to be a more general property of domains formed by membrane-proximal perilipins (Fig. S1B).

**Figure 3.**
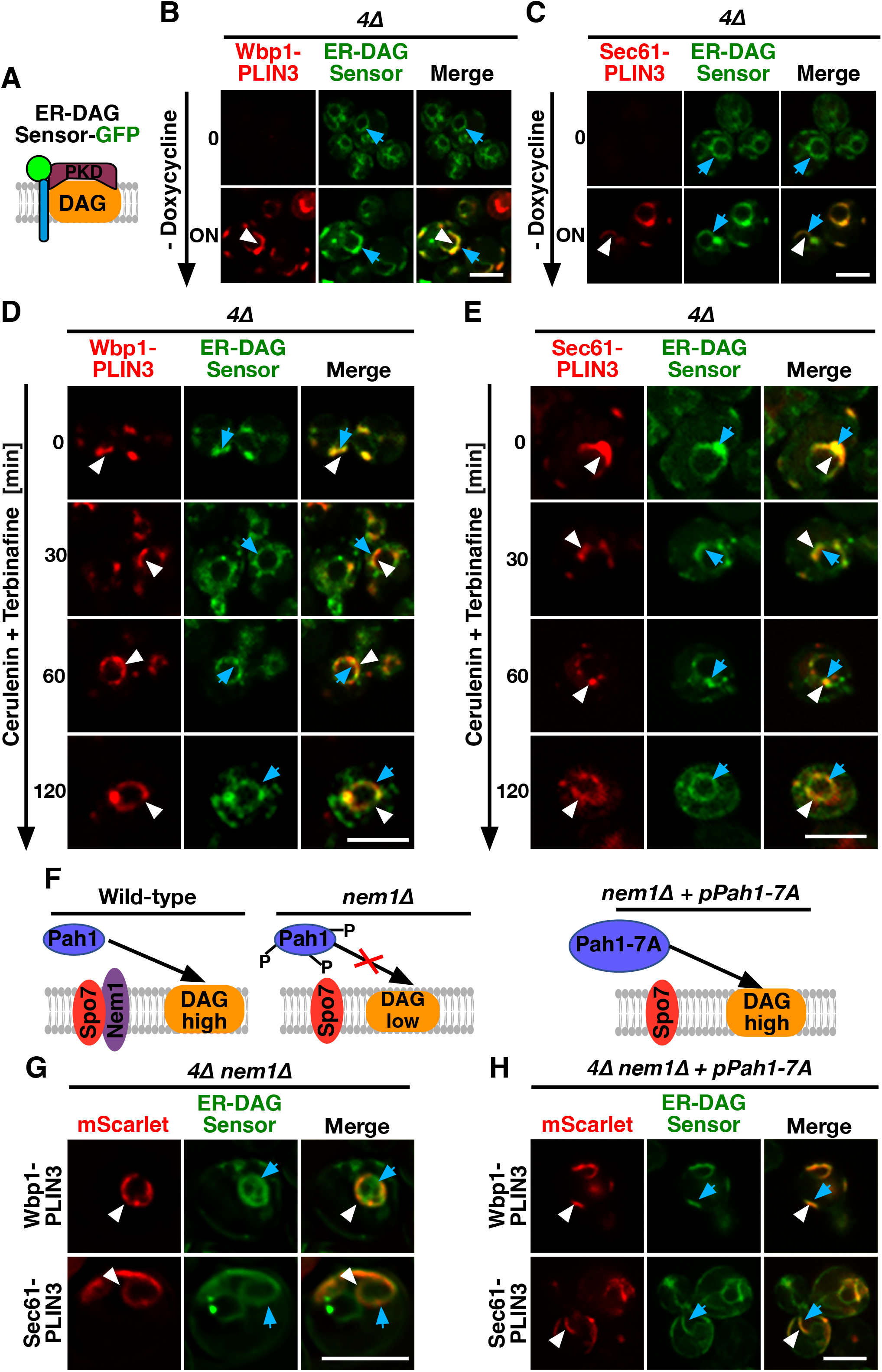
PLIN-induced ER domains colocalize with an ER-DAG sensor and are dependent on DAG formation. A) Schematic representation of the ER-DAG sensor. The ER-DAG sensor used here is composed of the DAG-binding tandem C1 domains of Protein Kinase D (C1a/b-PKD; lila box) fused to GFP (green circle), which in turn is fused to the transmembrane domain of Ubc6 (blue cylinder), a tail-anchored ER protein. DAG accumulation is shown in orange. B, C) Expression of the membrane-proximal PLIN3 induces colocalization of the ER-DAG sensor. Cells coexpressing inducible versions of either Wbp1-mScarlet-PLIN3 (B), or Sec61-mScarlet-PLIN3 (C) with a constitutively expressed GFP-tagged ER-DAG sensor, were imaged after growth in the presence (time 0) or absence of doxycycline (overnight, ON) to induce expression of the PLIN3 reporter. ER domains formed by membrane-proximal PLIN3 are indicated by white arrowheads, the localization of the ER-DAG sensor is indicated by blue arrows. Scale bar, 5 μm. D, E) PLIN3 membrane domains transform upon inhibition of lipid synthesis. Cells coexpressing the indicated mScarlet-tagged membrane-proximal PLIN3 together with the GFP-tagged ER-DAG sensor were treated with cerulenin (10 μg/ml) and terbinafine (30 μg/ml) for the indicated period of time and the distribution of the marker proteins was analyzed by confocal microscopy. Localization of the membrane-proximal PLIN3 is indicated by white arrowheads, that of the ER-DAG sensor by blue arrows. Scale bar, 5 μm. F-H) Formation of PLIN3 membrane domains depends on DAG synthesis. Schematic representation of the activation of the phosphatidate phosphatase Pah1 through dephosphorylation by the ER membrane embedded Nem1/Spo7 phosphatase complex. Dephosphorylation of Pah1 in wild-type cells promotes DAG production, whereas lack of Nem1 results in low DAG levels (F). ER domain formation was analyzed in mutants lacking the activator of Pah1, Nem1 (*are1*Δ *are2*Δ *dga1*Δ *lro1*Δ *nem1*Δ, panel G) and in *nem1*Δ mutant cells expressing a constitutive active version of Pah1, Pah1-7A (panel H). Cells co-expressing the membrane-proximal PLIN3 together with the ER-DAG sensor were imaged by confocal microscopy. Localization of the membrane-proximal PLIN3 is indicated by white arrowheads, that of the ER-DAG sensor by blue arrows. Scale bar, 5 μm.

Consistent with an important function of lipids in formation of the PLIN3 domains, simultaneously blocking the synthesis of fatty acids, by treating cells with cerulenin, and that of sterols, with terbinafine, resulted in a rapid dispersal of the PLIN3 domains and a more uniform ER localization of Wbp1-mScarlet-PLIN3 and Sec61-mScarlet-PLIN3 (Omura, 1981; Petranyi et al., 1984) (Fig. 3D, E). In wild-type cells, this block in lipid synthesis induces the mobilization of neutral lipids, TAG and STE, and hence turnover of LDs (Köffel et al., 2005). In the quadruple mutant used here, this block in lipid synthesis is likely to decrease DAG levels, resulting in a more homogenous distribution of the ER-anchored DAG sensor. Thus, DAG levels appear to be important for the formation of the PLIN3 domains as they rapidly dissolve upon depletion of DAG.

To further test whether elevated levels of DAG are required for the formation of PLIN3 membrane domains, we examined the localization of membrane-anchored PLIN3 in cells lacking Nem1. Nem1 is part of the heteromeric Nem1/Spo7 phosphatase complex, which activates the phosphatidate phosphatase Pah1, the key enzyme that converts phosphatidic acid (PA) into DAG (Kwiatek et al., 2020) (Fig. 3F). In the absence of Nem1, domain formation in the quadruple mutant was abrogated, indicating that DAG is required for the formation of membrane domains by PLIN3 (Fig. 3G). Mutant cells lacking Nem1 are known to have elevated levels of PA and they show proliferation of the nuclear ER, which explains the aberrant ER morphology observed in some of the cells expressing the membrane-anchored PLIN3 (Siniossoglou et al., 1998; Santos-Rosa et al., 2005). Formation of PLIN3 membrane domains in the *nem1*Δ mutant background, however, was restored upon expression of a constitutive active version of Pah1, Pah1-7A, thus confirming that DAG levels are important for domain formation (O’Hara et al., 2006) (Fig. 3F, H). These data thus indicate that PLIN3 induces the formation of membrane domains enriched in DAG and that DAG is required for the formation of these domains. Thus, PLIN3 appears to exert membrane organizing properties by concentrating DAG within the ER membrane bilayer. These high concentrations of DAG might then be recognized by otherwise LD-localized proteins such as Erg6 and by the neutral lipid dyes.

### PLIN3 containing ER domains are enriched in LD biogenesis proteins

Considering that membrane proximal PLIN3 induces membrane domains that have properties of LDs and are labelled by LD-localized proteins, we wondered whether these domains are also recognized by ER-localized proteins that function in the biogenesis of LDs. We have previously shown that the synthesis of TAG occurs at ER subdomains containing the LD biogenesis proteins seipin (composed of Fld1/Sei1 and Ldb16 in yeast), the regulators of Pah1 activity Nem1/Spo7, and Pex30, a reticulon homology domain containing protein that has membrane tubulating activity and is required for proper LD formation (Joshi et al., 2018; Wang et al., 2018; Choudhary et al., 2020) (Fig. 4A). We therefore tested whether these LD biogenesis proteins colocalize with PLIN3 crescents. Remarkably, most of the proteins tested, i.e., seipin, Ldb16, Nem1, Spo7, and Ldo16/45 colocalized with both Wbp1-GFP-PLIN3 and Sec61-GFP-PLIN3 within ER crescents, as shown by the line scans (Fig. 4B, C). Ldo16/45, are two splice-variants of lipid droplet organization protein (Ldo), which interact with the seipin complex and regulate LD metabolism (Teixeira et al., 2018; Eisenberg-Bord et al., 2018). Pex30, on the other hand, appeared to be enriched at one edge of the crescent-shaped domain, as evidenced by a yellow spot on the edge of the crescent in the merged image and the corresponding line scan. Taken together, these data indicate that either PLIN3 or DAG or their combination induce the localization of LD biogenesis proteins into ER subdomains.

**Figure 4.**
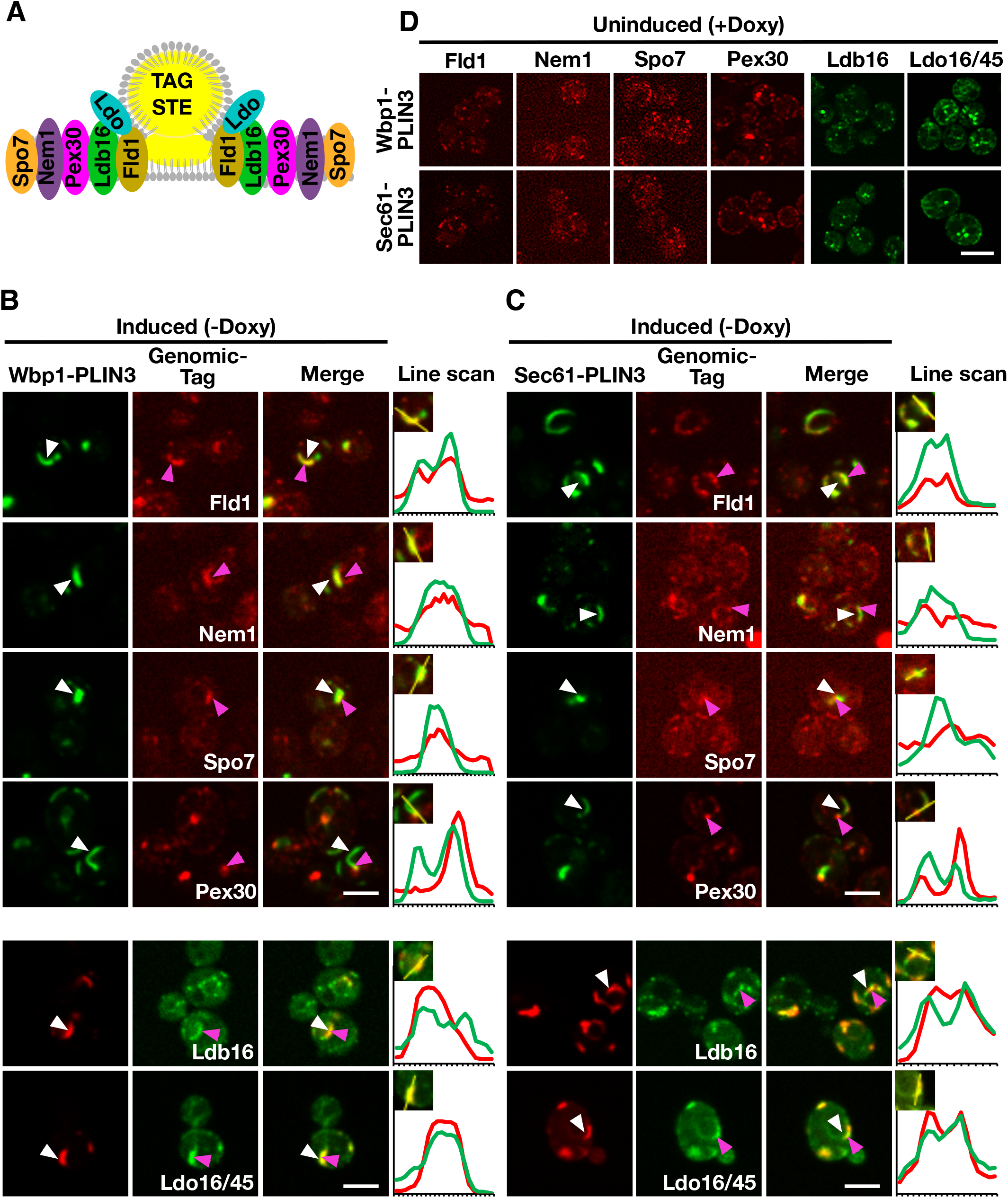
Membrane domains formed by PLIN3 are enriched in LD biogenesis proteins. A) Schematic representation of ER-localized LD biogenesis proteins. Seipin (Fld1), which in yeast forms a complex with Ldb16, colocalizes with the reticulon homology domain containing Pex30, the lipid droplet organizing proteins and promethin ortholog Ldo16/45, and the activator complex of the phosphatidate phosphatase Pah1, Nem1/Spo7, at the base of neutral lipid (TAG, STE, indicated in yellow) filled LDs. B-C) ER crescents formed by the membrane-anchored PLIN3 are enriched in the LD biogenesis proteins seipin (Fld1), Nem1, Spo7, Pex30, Ldb16, and Ldo16/45. Quadruple mutant cells co-expressing either GFP-(signal in green) or mScarlet-tagged (signal in red) versions of Wbp1-PLIN3 (B), or Sec61-PLIN3 (C), with the indicated genomically-tagged LD biogenesis protein were cultivated to early logarithmic growth phase in rich medium and the subcellular localization of the indicated fluorescent proteins was analyzed by confocal microscopy. Localization of the membrane-proximal PLIN3 is indicated by white arrowheads, that of the LD biogenesis protein by pink arrowheads. Regions selected for the line scans (yellow line) are shown in the insets and line scans are plotted to the right of the merged images. Scale bar, 5 μm. D) In the absence of expression of membrane-anchored PLIN3, LD biogenesis factors are localized to punctate structures, but do not form crescent-shaped membrane domains. Cells bearing plasmids containing Wbp1-mScarlet-PLIN3, or Sec61-mScarlet-PLIN3, were cultivated in the presence of doxycycline to repress expression of the membrane-anchored PLIN3 fusion proteins and the localization of the genomically-tagged versions of the LD biogenesis proteins was assessed by confocal microscopy. Scale bar, 5 μm.

### LD biogenesis proteins affect the number of ER crescents induced by membrane proximal PLIN3 and the distribution of the ER-DAG sensor

To test whether formation of the ER domain by the membrane-proximal PLIN3 and its colocalization with the ER-DAG sensor depends on LD biogenesis factors, we examined their localization in *4*Δ cells lacking seipin, Ldb16, or the Ldo16/45 proteins (i.e., a *ldo16*Δ *ldo45*Δ double mutant (Teixeira et al., 2018; Eisenberg-Bord et al., 2018). The membrane-proximal PLIN3 formed ER membrane domains and colocalized with the ER-DAG sensor in mutant cells lacking any one of these LD biogenesis factors, indicating that their function in LD biogenesis is not strictly required for the formation of the DAG-enriched ER domains (Fig. 5A, B). However, the size, shape, and number of the domains formed in the absence of these LD biogenesis proteins appear to be altered as the domains appeared to be smaller, less elongated, but more frequent. Indeed, quantification of the number of PLIN3 crescents and punctate structures formed indicates that *4*Δ mutant cells typically contain a single crescent-shaped domain, whereas the mutants in LD biogenesis proteins frequently contain two or even more of these structures (Fig. 5C, D). The LD biogenesis proteins also affected the distribution of the ER-DAG sensor, resulting in a more widespread localization of the sensor throughout the perinuclear ER (Fig. 5A-D). Taken together, these results indicate that LD biogenesis proteins affect the size, shape, and number of the PLIN-induced ER domains and the degree of localization of the ER-DAG sensor within these domains, indicating that these proteins affect the integrity, possibly the stability of the membrane domains formed by membrane-proximal PLIN3.

**Figure 5.**
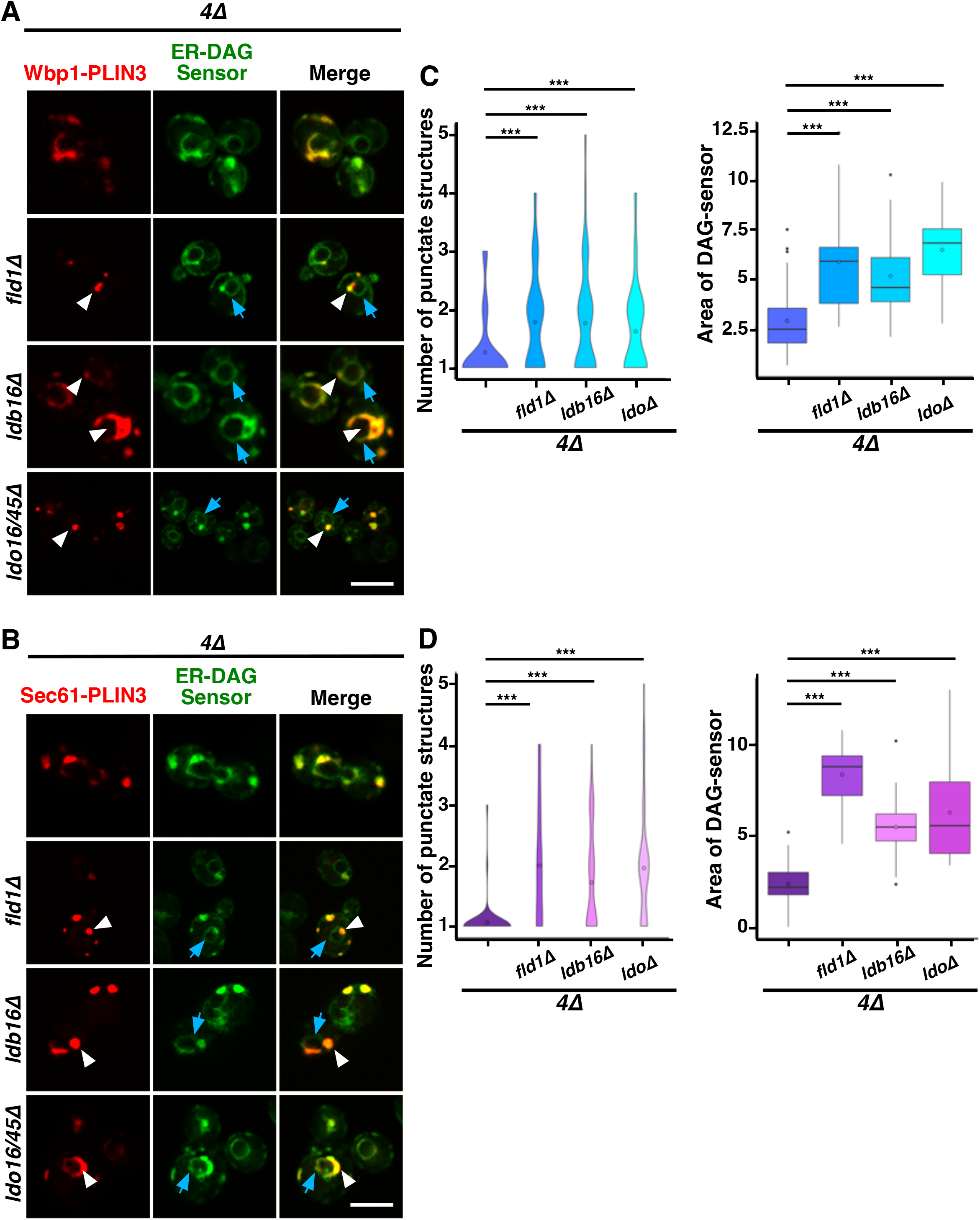
LD biogenesis proteins affect the size and number of ER domains formed by membrane-proximal PLIN3. A, B) Colocalization between membrane-proximal PLIN3 and the ER-DAG sensor was analyzed in *4*Δ cells (*are1*Δ *are2*Δ *dga1*Δ *lro1*Δ) lacking one of the indicated LD biogenesis genes (*fld1*Δ, or *ldb16*Δ, or *ldo16/45*Δ). Localization of the membrane-proximal PLIN3 fused to mScarlet and Wbp1-PLIN3 (panel A) or Sec61-PLIN3 (panel B) is indicated by white arrowheads, that of the ER-DAG sensor by blue arrows. Scale bar, 5 μm. C, D) Quantification of the average number of PLIN3 membrane domains per cell and the relative area occupied by the ER-DAG sensor in *4*Δ mutant cells lacking LD biogenesis proteins. The average number of PLIN3 membrane domains and the relative area occupied by the ER-DAG sensor was quantified manually in N> 50 cells and plotted as violin and box plot, respectively. The median was calculated by counting the number of punctate and measuring the area occupied by the ER-DAG sensor. The significance was assessed using a Wilcoxon rank-sum test with continuity correction. *\*\*\*P*<0.001.

### ER membrane domain formation by PLIN3 requires at least two of the three protein domains present in PLIN3

Given that the membrane-anchored fusion proteins containing a full-length PLIN3 induce formation of LD-like membrane-domains enriched in DAG, LD proteins, and LD biogenesis factors, when expressed in cells that are unable to generate storage lipids and LDs, we wondered whether this membrane domain-forming activity could be attributed to a particular protein domain within PLIN3. Most of the perilipin family members, including PLIN3, are composed of three distinct protein domains, an N-terminal PAT domain of unknown function, a central seven-fold 11-mer repeat domain composed of amphipathic helices, and a C-terminal 4-helix bundle domain (Kimmel & Sztalryd, 2016; Sztalryd & Brasaemle, 2017; Itabe et al., 2017) (Fig. 6A). To test whether any one of these three protein domains is sufficient for the induction of ER membrane domains, we generated constructs containing each one of the PLIN3 protein domains fused to the membrane anchor, i.e., Wbp1-mScarlet or Sec61-mScarlet, and tested whether these single-domain fusions would be sufficient to induce ER membrane domain formation in a quadruple mutant background. Western blot analysis of proteins extracted from quadruple mutant cells expressing these single PLIN3 domain fusion proteins indicate that all the fusion constructs were stably expressed (Fig. 6B). However, none of these single-domain fusions was able to induce the formation of membrane domains as both the mScarlet fusion reporter and the ER-DAG sensor exhibited uniform circular staining of the perinuclear ER (Fig. 6C, D). We thus generated all possible combinations of two-protein domain fusions, i.e., a PAT+11-mer repeat, PAT+4-helix bundle, and 11-mer+4-helix bundle combination. Remarkably each one of these double-protein domain fusions induced the formation of ER crescents and colocalized with the ER-DAG sensor (Fig. 6C, D). In addition, these double protein domain fusions also induced clustering of seipin, when expressed in a *4*Δ mutant background lacking LDs (Fig. S2). These results suggest that the 3 individual protein domains within PLIN3 exert both redundant and synergistic, or possibly sequential, functions in ER crescent formation, DAG accumulation, and clustering of LD biogenesis proteins. Thus, a single PLIN3 protein domain is not sufficient to induce ER crescent formation, but the combination of any two of these three PLIN3 protein domains is proficient in formation of ER crescents and their colocalization with the ER-DAG sensor.

**Figure 6.**
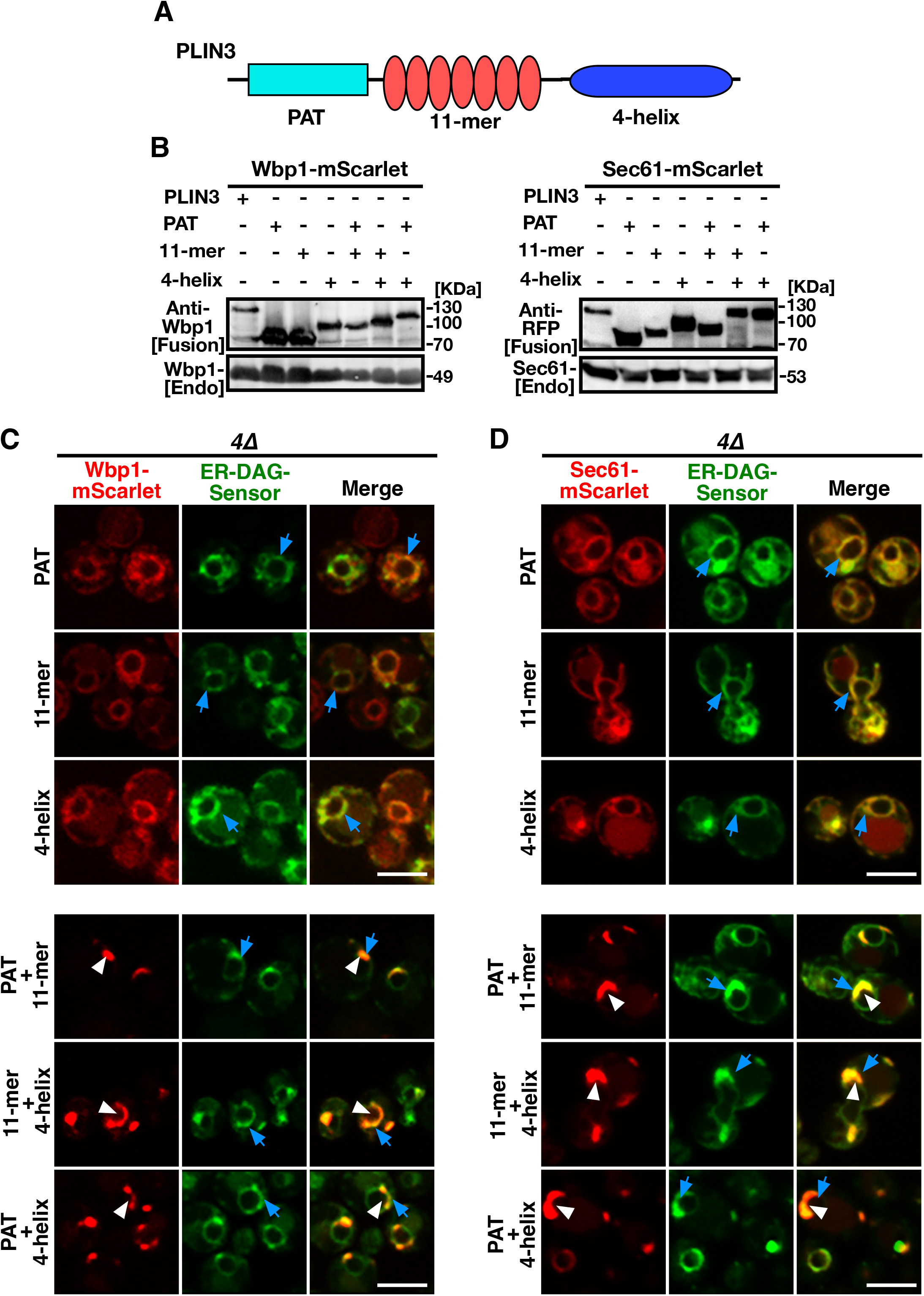
Protein domains within PLIN3 exert synergistic functions in ER membrane domain formation and clustering of DAG. A) Schematic representation of the domain structure of PLIN3. The three-domain architecture of PLIN3 is illustrated: N-terminal PAT domain (blue rectangle, amino acids 1-99), the repetition of 11-mer amphipathic helices (red ovals, amino acids 96-192), and the C-terminal 4-helix bundle (dark blue ellipse, amino acids 192-434). B) Western blot analysis of membrane-anchored PLIN3 protein domains. Proteins from cells expressing fusions of Wbp1 or Sec61 with either full-length PLIN3 or the indicated single- or double-PLIN3 protein domains were extracted, separated by SDS-PAGE, and probed for the presence of the fusion proteins [Fusion] or for endogenous native Wbp1 or Sec61 [Endo]. C, D) Any two of the three protein domains of PLIN3 are required for ER membrane domain formation. *4*Δ cells (*are1*Δ *are2*Δ *dga1*Δ *lro1*Δ) co-expressing the ER-DAG sensor together with single protein domains of PLIN3, i.e., PAT, 11-mer repeat, or 4-helix bundle, or two-domain fusions, i.e., PAT+11-mer, PAT+4-helix bundle, or 11-mer+4-helix bundle, fused to either Wbp1-mScarlet (C) or Sec61-mScarlet (D) were analyzed by confocal microscopy. Localization of the membrane-proximal PLIN3 domain in crescent shaped ER domains is indicated by white arrowheads, that of the ER-DAG sensor by blue arrows. Scale bar, 5 μm.

When expressed in wild-type cells, the single PLIN3 protein domain fusions localized to the ER and did not colocalize with LDs (Fig. S3). On the other hand, the PLIN3 two-domain fusions colocalized with LDs, indicating that crescent-shaped ER domain formation in cells lacking LDs correlates with the propensity of these domains to bind LDs and to induce a rearrangement of the ER around LDs (Khaddaj et al., 2022) (Fig. S3).

### PLIN3 binds to liposomes containing DAG

Given that membrane-proximal PLIN3 induce the formation of crescent-shaped ER membrane domains that are enriched in the ER-DAG sensor, we wondered whether PLIN3 might directly bind and cluster DAG within the ER membrane. To test this, we expressed a polyhistidine-tagged version of PLIN3 in bacteria and purified the protein by affinity chromatography. We then assessed whether the purified PLIN3 could bind DAG *in vitro* using microscale thermophoresis (Wienken et al., 2010; Seidel et al., 2013). When incubated with soluble short chain DAG analog, C8:0-DAG, PLIN3 bound this lipid with an apparent dissociation constant *K_D_* of 0.22 μM (Fig. 7A). To test whether PLIN3 would recognize membrane embedded DAG, we tested binding of PLIN3 to liposomes having a lipid composition similar to that of the ER membrane. PLIN3 bound to these ER liposomes with a *K_D_* of 409 μM, and this binding affinity was increased by more than 6-fold when 5 mol% long chain C16:0-DAG was present (Fig. 7B, C). These results indicate that PLIN3 can bind DAG and that the presence of DAG within a membrane enhances the binding affinity of PLIN3 towards the membrane.

**Figure 7.**
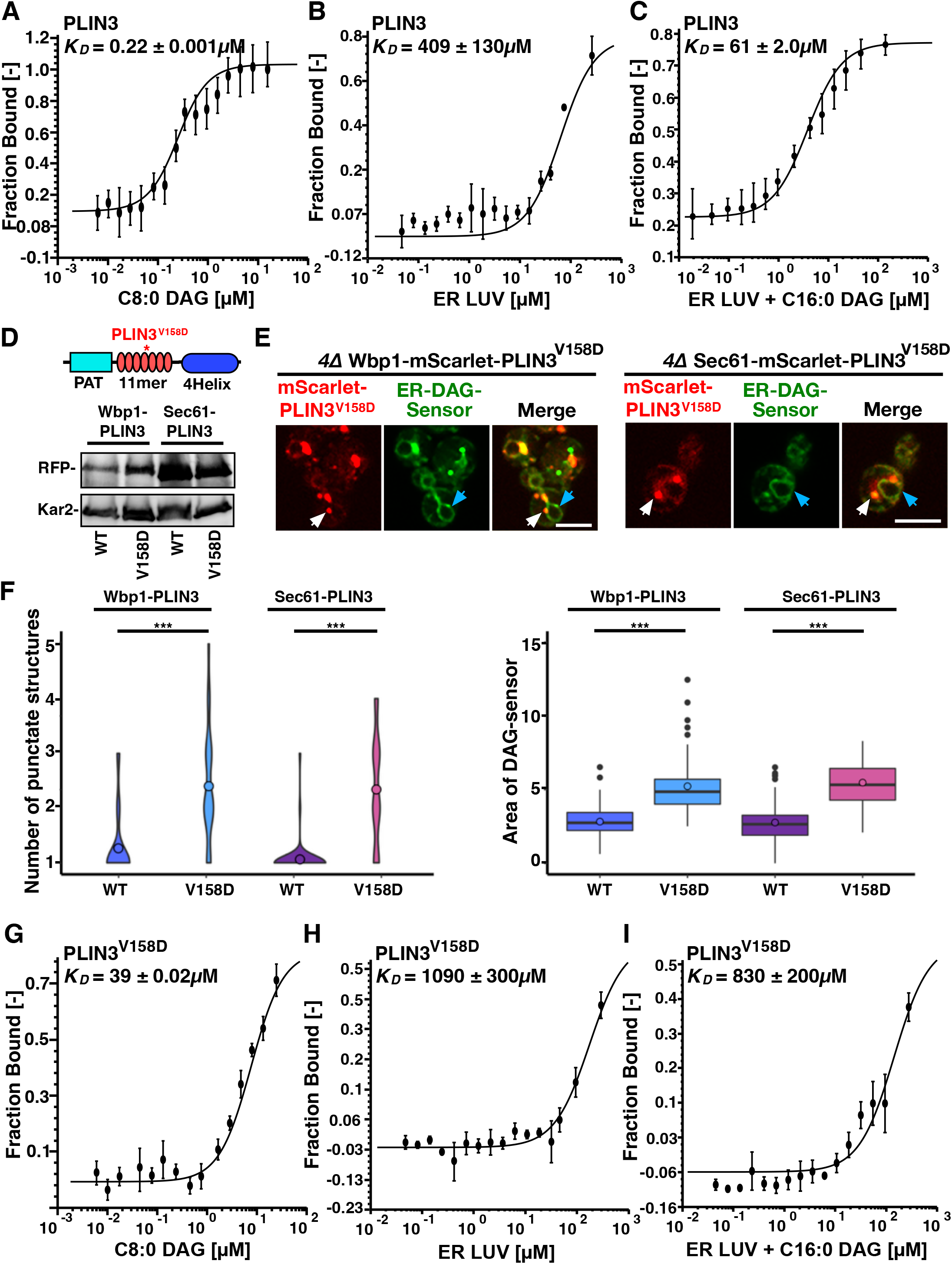
PLIN3 binds DAG-containing liposomes *in vitro*. A) Purified PLIN3 binds short chain DAG *in vitro*. An N-terminal polyhistidine-tagged version of PLIN3 was expressed in bacteria and purified by Ni^2+^-affinity chromatography. The protein was fluorescently labelled and binding of short chain DAG (C8:0-DAG) was measured by microscale thermophoresis. The fraction of the ligand-bound protein is plotted against the concentration of DAG and the deduced dissociation constant (*K_D_*) is indicated. Values represent mean ± S.D. of three independent determinations. B, C) PLIN3 displays increased binding to liposomes containing DAG. Purified and fluorescently labelled PLIN3 was incubated with liposomes lacking (panel B) or containing 5 mol% C16:0-DAG (1,2-dipalmitoyl-sn-glycerol, panel C) and the binding affinity was determined by microscale thermophoresis. The deduced dissociation constants (*K_D_*) are indicated. Values represent mean ± S.D. of three independent determinations. D) A mutant version of PLIN3, which bears a point mutation in the amphipathic helix repeats encoded by the 11-mer repeats, PLIN3^V158D^, is stably expressed. A schematic drawing of the structure of PLIN3 and the position of the point mutation is shown on top. Western blot analysis of cells expressing fusions between Wbp1 or Sec61 and either wild-type (wt) or the V158D mutant version of PLIN3. Kar2 is shown as a loading control. E) Expression of the point mutant version of membrane-anchored PLIN3 affect the number of membrane domains that are being formed and the relative distribution of the ER-DAG sensor. Cells co-expressing membrane-anchored versions of PLIN3^V158D^ with the ER-DAG sensor were cultivated and analyzed by confocal microscopy. Localization of the membrane-proximal PLIN3^V158D^ domain is indicated by white arrows, that of the ER-DAG sensor by blue arrows. Scale bar, 5 μm. F) The V158D mutant version of the membrane-anchored PLIN3 affects the numbers of punctate structures and the localization of the ER-DAG sensor. The number of punctuate structured formed by the wild-type and the V158D mutant version of the membrane-anchored PLIN3 is plotted as is the relative area of the ER-DAG sensor. The average number of PLIN3 membrane domains and the relative area occupied by the ER-DAG sensor was quantified manually in N>50 cells and plotted as violin and box plot, respectively. The median was calculated by counting the number of punctate and measuring the area occupied by the ER-DAG sensor. The significance was assessed using a Wilcoxon rank-sum test with continuity correction. *\*\*\*P*<0.001. G) The PLIN3^V158D^ point mutant has reduced affinity to short chain DAG. Purified PLIN3^V158D^ was fluorescently labelled and binding of short chain DAG (C8:0-DAG) was measured by microscale thermophoresis. The deduced dissociation constant (*K_D_*) is indicated. Values represent mean ± S.D. of three independent determinations. H, I) PLIN3^V158D^ has reduced binding affinity towards liposomes containing DAG. PLIN3^V158D^ was incubated with liposomes lacking (panel H) or containing 5 mol% C16:0-DAG (1,2-dipalmitoyl-sn-glycerol, panel I) and the binding affinity was determined by microscale thermophoresis. The deduced dissociation constants (*K_D_*) are indicated. Values represent mean ± S.D. of three independent determinations.

To test the specificity of DAG binding, we used a version of PLIN3, which bears a point mutation in the fifth of a total of seven amphipathic helices. This PLIN3^V158D^ mutant version has previously been shown to affect LD targeting of the soluble PLIN3 protein when expressed in yeast cells (Rowe et al., 2016). To test whether this PLIN3 mutant version retains the capacity to induce membrane domains, we first expressed a membrane-anchored versions of PLIN3^V158D^ in the quadruple mutant co-expressing the ER-DAG sensor. The membrane-anchored PLIN3^V158D^ mutant versions were stably expressed as revealed by Western blot analysis (Fig. 7D). The PLIN3^V158D^ mutant versions still induced the formation of punctate and crescent-shaped ER membrane domains. These domains, however, did not colocalize with the ER-DAG sensor to the same degree as observed with wild-type PLIN3, suggesting that the propensity to form DAG-enriched membrane domains is reduced by the point mutation in PLIN3 (Fig. 7E). The membrane domains formed by the mutant version of PLIN3, were more numerous, and smaller compared to those formed by wild-type PLIN3 (Fig. 7F, 5E, 1F, 1G). In addition, the ER-DAG sensor was spread over a larger area of the ER and was not as strongly recruited to the PLIN3^V158D^-induced domains, as compared to cells expressing wild-type PLIN3 (Fig. 7F). This increase in the number of ER membrane domains, and wider dispersion of the ER-DAG sensor, observed in cells expressing the mutant version of PLIN3 is thus similar to the changes observed in mutant cells lacking individual LD biogenesis proteins (Fig. 5). These results indicate that a mutation in the amphipathic helices of PLIN3, which is known to affect targeting of the soluble protein to LDs, also affects the number and size of the ER membrane domains and the recruitment of the ER-DAG sensor to these domains.

To test whether the point mutant version of PLIN3 directly affects binding of DAG, the mutant protein was expressed in bacteria and was affinity purified. Lipid binding assays revealed that the PLIN3^V158D^ mutant version had a greatly reduced affinity towards short chain C8:0-DAG (*K_D_* of 39 μM), compared to the wild-type protein (*K_D_* of 0.22 μM), revealing a 177-fold reduced binding affinity of the mutant protein compared to wild-type PLIN3 (Fig. 7G). In addition, the PLIN3^V158D^ mutant version only poorly bound to liposomes lacking C16:0-DAG (*K_D_* of 1090 μM), or to liposomes containing C16:0-DAG (*K_D_* of 830 μM) (Fig. 7H, I). Given that the mutant version of PLIN3 exhibits a 177-fold decreased binding to short chain DAG and a 13-fold reduced binding to ER liposomes containing 5 mol% DAG when compared to the wild-type PLIN3, the data indicate that formation of ER membrane domains by the membrane-anchored PLIN3 and its propensity to recruit the ER-DAG sensor correlate with the affinity of the protein to bind short chain DAG and to bind to liposomes containing low concentrations of DAG. Taken together, these results indicate that wild-type PLIN3 exerts binding specificity to membranes containing DAG, and that this specificity requires the function of all seven of the amphipathic helices within the 11-mer repeats of PLIN3, because a single point mutation, V158D, in the fifth 11-mer repeats greatly reduces liposome binding.

## Discussion

In this study, we employed membrane-proximal versions of PLIN3 to characterize the function and molecular properties of perilipins. We have previously shown that expression of membrane-anchored PLIN3 in cells containing LDs induces rearrangement of the ER membrane around LDs. This wrapping of LDs by the ER membrane is so tight that, at the resolution of the light microscope, the PLIN reporters appear to colocalize with genuine LD marker proteins (Khaddaj et al., 2022) (Fig. 1C). Here, we now expressed membrane-anchored PLIN reporters in cells lacking LDs. Under these conditions, these reporters (PLIN3 and PLIN1) concentrate in crescent-shaped ER domains (Fig. 1D, E, S1). These ER domains bear hallmarks of LDs as they are recognized by LD-localized proteins (Erg6, Tgl1, Tgl4, Ayr1) as well as fluorescent neutral lipid-specific dyes (BODIPY, Nile Red) (Fig. 2). Upon induction of TAG synthesis, these crescent-shaped ER domains transform into smaller circular shaped LDs indicating that these domains can be resolved and hence are not caused by an irreversible aggregation of PLINs (Fig. 2). Their stability appears to depend on lipids, because they disperse when cells are treated with inhibitors that block lipid synthesis (cerulenin and terbinafine). Moreover, these ER domains colocalize with an ER-anchored sensor of DAG, suggesting that they contain elevated levels of DAG, the lipid precursor for synthesis of the major storage lipid, TAG. DAG is required for the formation of these ER domains, because they do not form in mutant cells lacking Nem1, a component of the Nem1/Spo7 phosphatase complex that is required for activation of Pah1, the phosphatidate phosphatase that converts PA into DAG (Fig. 3).

DAG has previously been shown to be required for LD formation, independent of its metabolic function as precursor to TAG formation (Adeyo et al., 2011; Choudhary et al., 2018; Zoni et al., 2021a). Thus, the formation of these ER membrane domains by PLIN3 requires DAG and results in the concentration of DAG within the domains, suggesting that PLIN3 and DAG cooperate in domain formation and that this cooperation could be important for the role that DAG plays in LD formation. Interestingly, in wild-type cells, DAG is also enriched at sites of LD biogenesis, which form upon the colocalization of seipin with Nem1 and recruitment of one of TAG biosynthetic enzymes (Choudhary et al., 2020; Choudhary & Schneiter, 2021). Moreover, molecular dynamics simulations indicate that the seipin complex in the ER concentrates both TAG and DAG within its membrane-embedded inner ring structure (Prasanna et al., 2021; Zoni et al., 2021b; Klug et al., 2021). Thus, the crescent-shaped ER domains formed by membrane-anchored PLIN3 might represent an enlarged and possibly more stable form of the microdomains that naturally occur at early stages of LD biogenesis. This points to a function of the soluble PLIN3 in these early stages of LD biogenesis, where it may act more transiently to concentrate both DAG and LD biogenesis proteins (Skinner et al., 2009). Consistent with such a membrane organizing function of PLIN3, the ER domains formed by expression of membrane-anchored PLIN3, in cells lacking the capacity to form neutral lipids, are enriched in LD biogenesis proteins, including seipin, Ldb16, Pex30, the Ldo16/45 proteins, as well as the Nem1/Spo7 complex (Fig. 4). Unlike Nem1, however, these LD biogenesis factors are not required for formation of the PLIN3/DAG domains, but they nevertheless appear to affect the relative number and size of the domains that are being formed. While cells lacking the capacity to form neutral lipids typically show one large crescent-shaped PLIN3/DAG domain, these domains are frequently smaller but more numerous in mutants lacking LD biogenesis proteins (Fig. 5). While these domains still colocalize with the ER-DAG sensor, the sensor also stains a larger portion of the circular perinuclear ER, suggesting that membrane-anchored PLIN3 induced domain formation is affected in the absence of one of these LD biogenesis factors, possibly through the levels of DAG formation, its stability, or its distribution within the membrane.

To examine whether ER membrane domain formation could be attributed to one particular domain of PLIN3, we generated fusions between the membrane anchors, Wbp1 or Sec61, and the PAT, 11-mer repeat, or 4-helix bundle domain of PLIN3. These single-domain fusions displayed homogenous ER distribution as did the co-expressed ER-DAG sensor, indicating that membrane domain formation and DAG clustering cannot by assigned to a single domain of PLIN3. To test whether two of the three domains of PLIN3 would be sufficient for membrane domain formation and DAG clustering, we generated all possible two domain fusion proteins, i.e., PAT+11-mer repeat, PAT+4-helix bundle, and 11-mer repeat+4-helix bundle. Remarkably, al Thel three of these two domain fusions formed domains in the ER and affected the distribution of the ER-DAG sensor, suggesting that the three domains of PLIN3 function synergistically in domain formation, and that at least two of these domains are required for domain formation (Fig. 6).

Consistent with a possible direct function of PLIN3 in the formation of DAG-enriched membrane domains, PLIN3 binds short chain DAG *in vitro* with low micromolar affinity and shows a 6.7-fold higher binding affinity to liposomes containing long-chain DAG compared to liposomes lacking DAG (Fig. 7). Binding to short chain DAG or to liposomes containing DAG is impaired in a mutant version of PLIN3 bearing a point mutation that affects the amphipathic nature of helices within the 11-mer repeat domain. While the membrane anchored PLIN3^V158D^ mutant still appears to form punctate domains in the ER, these domains are smaller and more numerous, and they fail to recruit the ER-DAG sensor with the same propensity as observed for wild-type PLIN3. These observations suggest that the amphipathic nature of the helices of the 11-mer repeats of PLIN3 is important for DAG binding *in vitro* and for concentration of DAG in the ER membrane *in vivo*, as monitored by the ER-DAG sensor. Binding of PLIN3 to membranes enriched in DAG is consistent with previous observations that treatment of preadipocytes with a membrane-permeable DAG analog promotes recruitment of PLIN3, PLIN4, and PLIN5 to the ER, and that inhibition of the DAG lipase enhances recruitment of PLIN3 to the LD surface (Skinner et al., 2009).

PLIN3/TIP47 has previously been shown to promote the transformation of liposomes into lipid discs in *vitro* (Bulankina et al., 2009). This membrane remodeling function was attributed to the C-terminal 4-helix bundle of PLIN3, which shows structural homology to the N-terminal domain of apolipoprotein E (ApoE), a protein known to promote formation of small lipidic discs (Wilson et al., 1991; Saito et al., 2001; Hickenbottom et al., 2004). In addition, PLIN4, which lacks the PAT domain, forms stable lateral arrangements of amphipathic helices on a membrane surface, suggestive of a membrane coat (Giménez-Andrés et al., 2021; Copic et al., 2018). These independent observations are consistent with a more general membrane-organizing property of the native soluble PLINs, which is likely enhanced in the membrane-anchored PLIN fusions that we employed and characterized in this study.

Taken together, these data support a function of PLINs in organizing membrane properties. By binding to and thereby possibly concentrating DAG, together with LD biogenesis proteins, such as seipin at ER domains, perilipins could promote early stages of LD biogenesis. These functions of the naturally occurring soluble PLINs are in part deduced from our observations of the domain forming properties of membrane-anchored PLINs. Such a membrane proximal positioning of PLINs is likely to increase the natural propensity of PLINs to bind specific lipids and thereby induce the formation of membrane domains. It is interesting to note that membrane domain formation by PLINs is even sufficient to promote recruitment of proteins that are typically found on mature LDs, such as Erg6 or the lipases Tgl1 and Tgl4. Moreover, these domains are specifically stained by fluorescent dyes such as Nile Red or BODIPY and thus appear to mimic to some extent the biophysical properties of LDs. Thus, membrane binding by PLINs appears to be sufficient to impose LD-like properties onto the ER bilayer membrane. Further studies are now required to understand the functions of the different domains of PLINs in their synergistic or possibly sequential action in binding specific lipids, such as DAG, and in forming lateral membrane domains.

## Acknowledgements

We thank all members of the lab for support, advice, and discussions; Stéphanie Cottier for comments on the manuscript; Vineet Choudhary for helpful discussions; and the Light Microscopy and Image Analysis core facility of the University of Fribourg for support and assistance in this work. This work is supported by the Swiss National Science Foundation (31003A_17303 and 310030_207870).

## Materials and Methods

### Culture media and growth conditions

Yeast strains were cultured either in YP medium [1% bacto yeast extract, 2% bacto peptone (US Biological, Swampscott, MA)] or in synthetic complete (SC) medium [0.67% yeast nitrogen base without amino acids (US Biological), 0.73 g/l amino acids)], containing either 2% glucose or 2% galactose. Fatty acid-supplemented medium contained 0.24% Tween 40 (Sigma-Aldrich, St Louis, MO) and 0.12% oleic acid (Carl Roth, Karlsruhe, Germany). Synthesis of fatty acids and sterols was inhibited by the addition of cerulenin (10 μg/ml; Enzo Life Sciences, Inc., Farmingdale, NY) and terbinafine (30 μg/ml; Sigma-Aldrich, St Louis, MO). Expression of the membrane-proximal PLIN3 constructs was repressed by cultivating cells in the presence of doxycycline (1 μg/ml; Sigma-Aldrich, St Louis, MO) and expression was induced by shifting cells to media lacking doxycycline, typically 14-16 h before imaging.

### Yeast strains and plasmids

Yeast strains and plasmids used in this study are listed in Table S1 and S2. Deletion mutants were generated either by mating or by PCR-based targeted homologous recombination to replace the ORF of the gene of interest with a deletion cassette followed by a marker rescue strategy (Gueldener et al., 2002). Chromosomally C-terminal–tagged versions of reporter proteins (Erg6-mCherry, Nem1-mScarlet, Pet10-mScarlet, Pex30-mScarlet, Fld1-mScarlet, Ldb16-GFP and Ldo16/45-GFP) were constructed using PCR-based homologous recombination (Longtine et al., 1998; Bähler et al., 1998).

Plasmid pCM189 containing the tetracycline-repressible *tetO*_7_ promoter (Garí et al., 1997) was used as a backbone to express the membrane-anchored PLIN fusion proteins as previously described (Khaddaj et al., 2022). The mutant version of PLIN3 that affects the hydrophobicity of the amphipathic helix, PLIN3^V158D^, was generated by PCR-mediated site directed mutagenesis and cloned as a fusion with Wbp1-mScarlet and Sec61-mScarlet into PCM189 to generate PCM189-Wbp1-mScarlet-PLIN3^V158D^ and PCM189-Sec61-mScarlet-PLIN3^V158D^.

PLIN3 domain truncations were generated by PCR and cloned as fusions with Wbp1-mScarlet and Sec61-mScarlet into PCM189.

To study the single domains of PLIN3, the PAT domain of PLIN3 (amino acids 1-99) was cloned in frame with Wbp1-mScarlet or Sec61-mScarlet into pCM189. The 11-mer repeat region covered amino acids 96-192, and the 4-helix bundle amino acids 193-434. The two domain fusions were generated by PCR ligation of the respective single domains. All constructs were verified by sequencing (Microsynth AG, Buchs, Switzerland).

### Western blot analysis

Cells were grown overnight in SC media and 3 OD_600_ units were harvested. Cells were incubated with 1.85 M NaOH, proteins were precipitated using 20% TCA, and resuspended into sample buffer (Horvath & Riezman, 1994). Proteins were separated by SDS-PAGE, blotted and membranes were incubated with primary antibodies against Wbp1 (M. Aebi, ETH Zurich, Switzerland, diluted at 1:2000), or Sec61 (R. Schekman, diluted at 1:2000). mScarlet-tagged fusion proteins were detected using a, RFP mouse monoclonal antibody (ChromoTek GmbH, Martinsried, Germany, diluted at 1:2000, #6g6-20). Primary antibodies were detected by using horseradish peroxidase (HRP)-conjugated secondary antibodies (goat anti-mouse IgG (H+L)-HRP conjugate (1:10,000, Bio-Rad #1706516), goat anti-rabbit IgG (H+L)-HPR conjugate (1:10,000, Bio-Rad #1706515). Western blots experiments were repeated at least two times with essentially similar results.

### LD staining and live cell microscopy

Yeast cells were grown to early logarithmic phase (~1 OD_600nm_), pelleted by centrifugation, and resuspended in a small volume of media. 3 μl of the cell suspension were mounted on a glass slide and covered with an agarose patch. To stain LDs with a lipophilic dye, cells expressing Wbp1-mScarlet-PLIN3, Sec61-mScarlet-PLIN3, Wbp1-GFP-PLIN3, or Sec61-GFP-PLIN3 were grown in SC media to early stationary phase. Cells were incubated with either BODIPY 493/503 (0.5 μg/ ml; Invitrogen, Waltham, MA, USA) or with Nile Red (10 μg/ ml; Sigma-Aldrich, St Louis, MO) for 5 min at room temperature, washed twice with PBS, and resuspended in fresh medium. Images of Nile Red stained cells were collected using a Leica TCS SP5 (Leica, Wetzlar, Germany) confocal microscope equipped with a 63x/1.20 HCX PL APO objective, and a LAS AF software. Analysis of BODIPY stained cells and localization of mCherry-, mScarlet- and GFP-tagged fusion proteins were performed using a Visitron spinning disc CSU-W1 set-up (Visitron Systems, Puchheim Germany), consisting of a Nikon Ti-E inverted microscope, equipped with a CSU-W1 spinning disk head (50-μm pinhole disk; Yokogawa, Tokyo, Japan), an Evolve 512 (Photometrics) EM-CCD camera, and a PLAN APO 100x NA 1.3 oil objective (Nikon). Images were treated using ImageJ software and then resized in Photoshop (Adobe, Mountain View). Microscopic experiments were performed three times with essentially similar results.

Colocalization was evaluated manually by scoring color overlap in 100 cells (% colabelling per cell). Box plots, violin plots and statistical analysis were performed with R software (http://www.rproject.org/), and the ggplot2 package (Wickham, 2016). The median is indicated in a box that represents the 25-75^th^ percentile range. The whiskers denote the largest and smallest values with 1.5x of the interquartile range from the hinges of the box. Outliers are presented by dots. In the violin plot, the median is represented in the center of the box. The length of the violin represents the interquartile range. A Wilcoxon rank-sum test and signed-ranked test with continuity correction were used to assess the significance of data.

### Expression and purification of PLIN3

DNA encoding wild-type and the mutant version of PLIN3 were cloned into BamHI and NdeI restriction sites of pET16b vector (Novagen, Merck, Kenilworth, NJ, USA) using Gibson Assembly Cloning Kit (New England Biolabs, Ipswich, MA, USA). Plasmids were transformed into *E. coli* BL21, and expression of the proteins containing an N-terminal polyhistidine tag was induced with IPTG (1 mM) for 3 h at 30°C. Cells were collected, lysed, and the soluble fraction of the lysate was incubated with Ni^2+^-nitrilotriacetic acid (NTA) beads (Qiagen, Hilden, Germany). Proteins were eluted with 300 mM imidazole in 60 mM NaH_2_PO_4_, 60 mM NaCl, pH 8.0. The eluted proteins were applied to Zeba spin desalting columns (Thermo Fisher Scientific, Waltham, MA, USA), and the buffer was exchanged to 60 mM NaH_2_PO_4_, 10 mM NaCl, pH 8.0. Protein concentration was determined by Lowry assay using Folin reagent and BSA as standard.

### Microscale thermophoresis (MST)

Binding of PLIN3 to short chain C8:0-DAG (1,2-dioctanoyl-sn-glycerol; Avanti Polar Lipids, Birmingham, AL, USA, #800800) and to liposomes was assessed by MST using a Monolith NT.115 system (Nanotemper Technologies, Munich, Germany). Proteins were labeled using the RED-tris-NTA His tag protein-labeling kit (Nanotemper Technologies). Labeled proteins were added to a serial dilution of unlabeled DAG or liposomes prepared in binding buffer (20 mM Tris, pH 7.5, 30 mM NaCl, and 0.05% Triton X-100). Samples were loaded into MST standard capillaries, and MST measurements were performed using 80% laser power setting. The dissociation constant *K_D_* was obtained by plotting the normalized fluorescence (Fnorm or the fraction bound) against the logarithm of ligand concentration. Experiments were performed in triplicates and data were fitted using the *K_D_* model of the MO.Affinity Analysis software (Nanotemper Technologies).

### Liposome preparation

Dry films containing the desired amount of phospholipids with or without 5 mol% 1,2-dipalmitoyl-sn-glycerol (C16:0-DAG; Avanti Polar Lipids, #800816) were prepared from stock solutions of lipids in chloroform using a rotary evaporator. Liposomes mimicking the phospholipid composition of the ER membrane contained 50.5 mol% DOPC, 20.5 mol% DOPE, 8 mol% DOPS, 5 mol% DOPA, 11 mol% soy PI and 5 mol% C16:0-DAG (Avanti Polar Lipids, DOPC #850375, DOPS #840035, DOPA #840875, soy PI #840044). The phospholipid film was resuspended in 1 ml liposome buffer (50 mM NaCl, 25 mM Tris, pH 7.4) at a concentration of 2 mM phospholipids. After ten cycles of freezing in liquid nitrogen and thawing in a water bath at 55°C, the resulting multilamellar liposome suspension was extruded nineteen times through a polycarbonate filter of 1 μm pore size to generate large unilamellar vesicles (LUVs).

## Source Data

Figure 6-Source Data container contains 4 files. The 3 .tif files show unmodified source data of the Western blots shown in Fig. 6B (Anti-Sec61, Anti-Rfp, and Anti-Wbp1). The Pdf file highlights the region of the blots that were assembled in Fig. 6B.

Figure 7-Source Data container contains 3 files. The 2 .tif files show unmodified source data of the Western blots shown in Fig. 7D (Anti-Kar2, and Anti-Rfp). The Pdf file highlights the region of the blots that were assembled in Fig. 7D.

## Supplementary Data

### Supplementary Figure Legends

**Figure S1.**
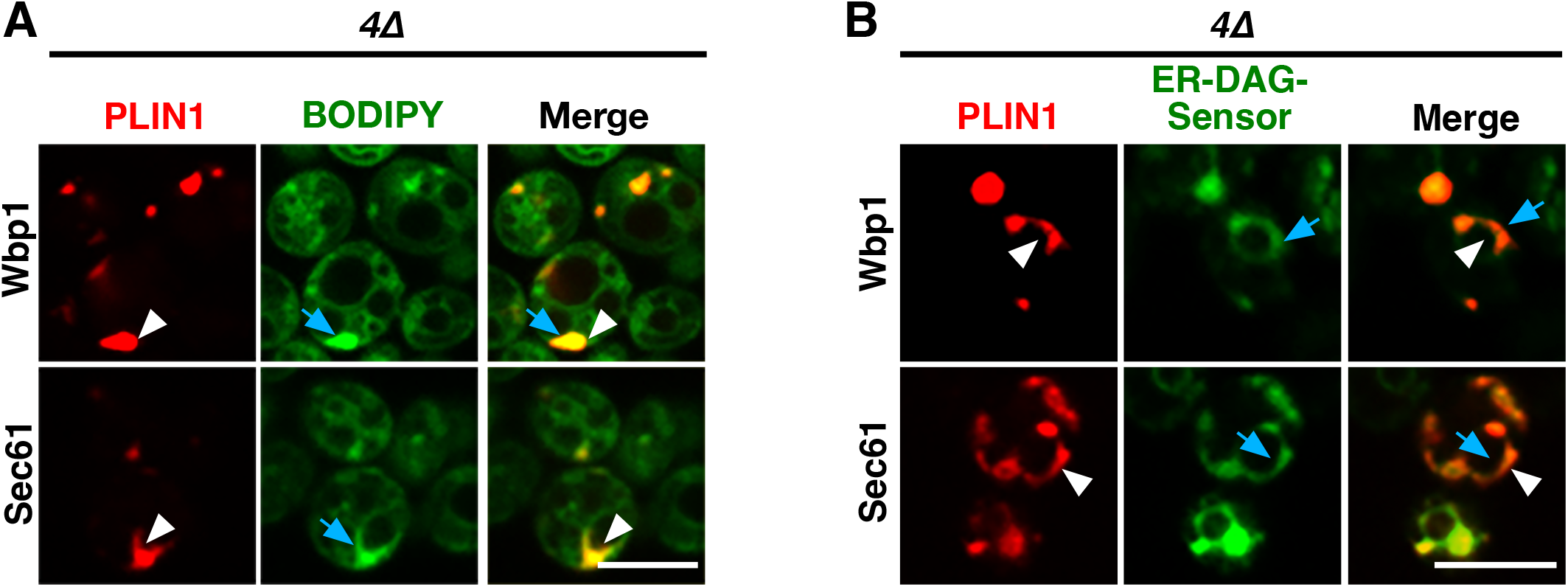
Fusion of PLIN1 to Wbp1 or Sec61 membrane anchors also induces formation of ER domains that colocalize with BODIPY or with the ER DAG-sensor. A) Quadruple mutant cells (*4*Δ; *are1*Δ *are2*Δ *dga1*Δ *lro1*Δ) expressing Wbp1-GFP-PLIN1 or Sec61-GFP-PLIN1 were stained with BODIPY and the distribution of the fluorescent-tagged proteins was analyzed by confocal microscopy. The ER domains formed by expression of the membrane-anchored PLIN1 are indicated by white arrowheads, co-staining by BODIPY is indicated by blue arrows. Scale bar, 5 μm. B) Quadruple mutant cells coexpressing Wbp1-mScarlet-PLIN1 or Sec61-mScarlet-PLIN1 with the GFP-tagged ER-DAG sensor were cultivated and their distribution was analyzed by confocal microscopy. ER domains formed by membrane-anchored PLIN1 are indicated by white arrowheads, that of the ER-DAG sensor by blue arrows. Scale bar, 5 μm.

**Figure S2.**
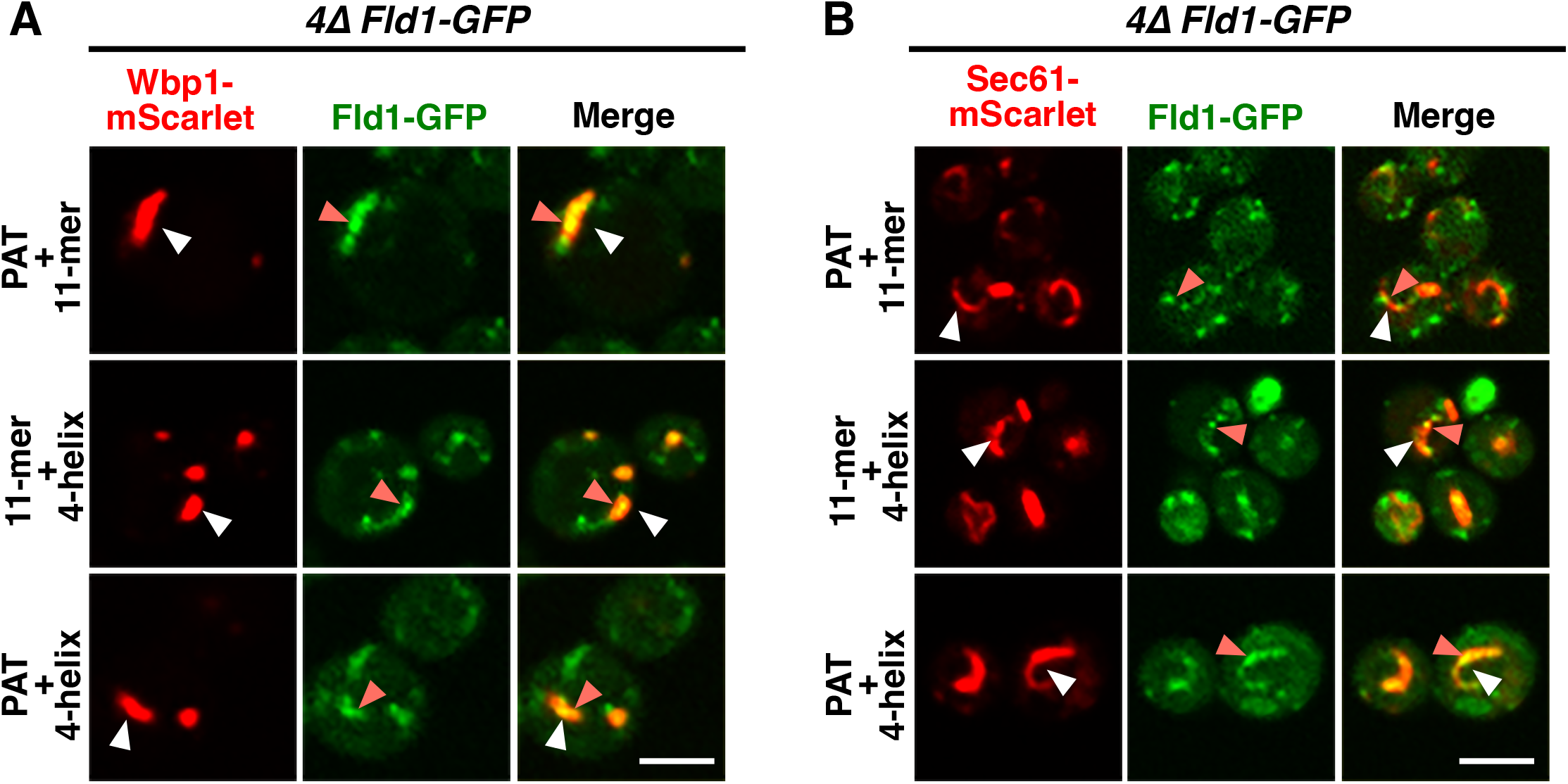
Seipin colocalizes with ER domains formed by membrane-anchored PLIN3 two-domain fusions. A, B) Quadruple mutant cells expressing GFP-tagged seipin (Fld1) and the indicated Wbp1-(panel A) or Sec61-(panel B) based PLIN3 double domain fusions were imaged by confocal microscopy. Membrane domains formed by the membrane-anchored PLIN double domain fusions are indicated by white arrowheads, foci formed by seipin are indicated by orange arrow heads. Scale bar, 5 μm.

**Figure S3.**
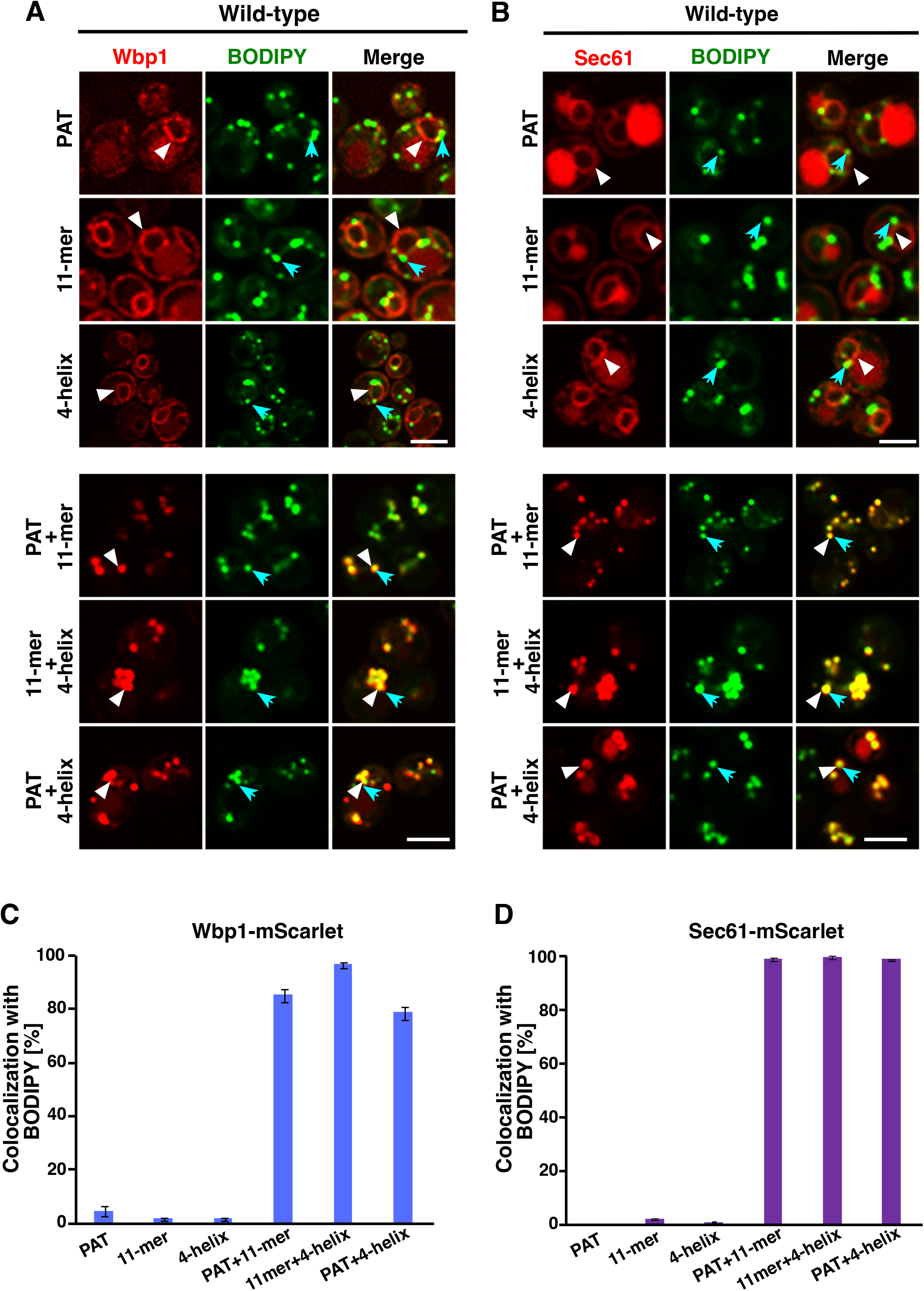
Membrane-anchored two PLIN3 protein domain fusions colocalize with LDs in wild-type cells. A, B) Wild-type cells expressing the indicated membrane-anchored PLIN3 single or double protein domain fusions were incubated with BODIPY to stain LDs and imaged by confocal microscopy. Localization of the membrane-anchored PLIN3 fusions is indicated by white arrowheads, BODIPY stained LDs are indicated by blue arrows. Scale bar, 5 μm. C, D) Quantification of colocalization between the membrane-anchored PLIN3 domain reporter and BODIPY stained LDs. Colocalization was scored manually, data represent mean ± S.D. of n=LDs cells.

### Supplementary Tables

**Table S1.**
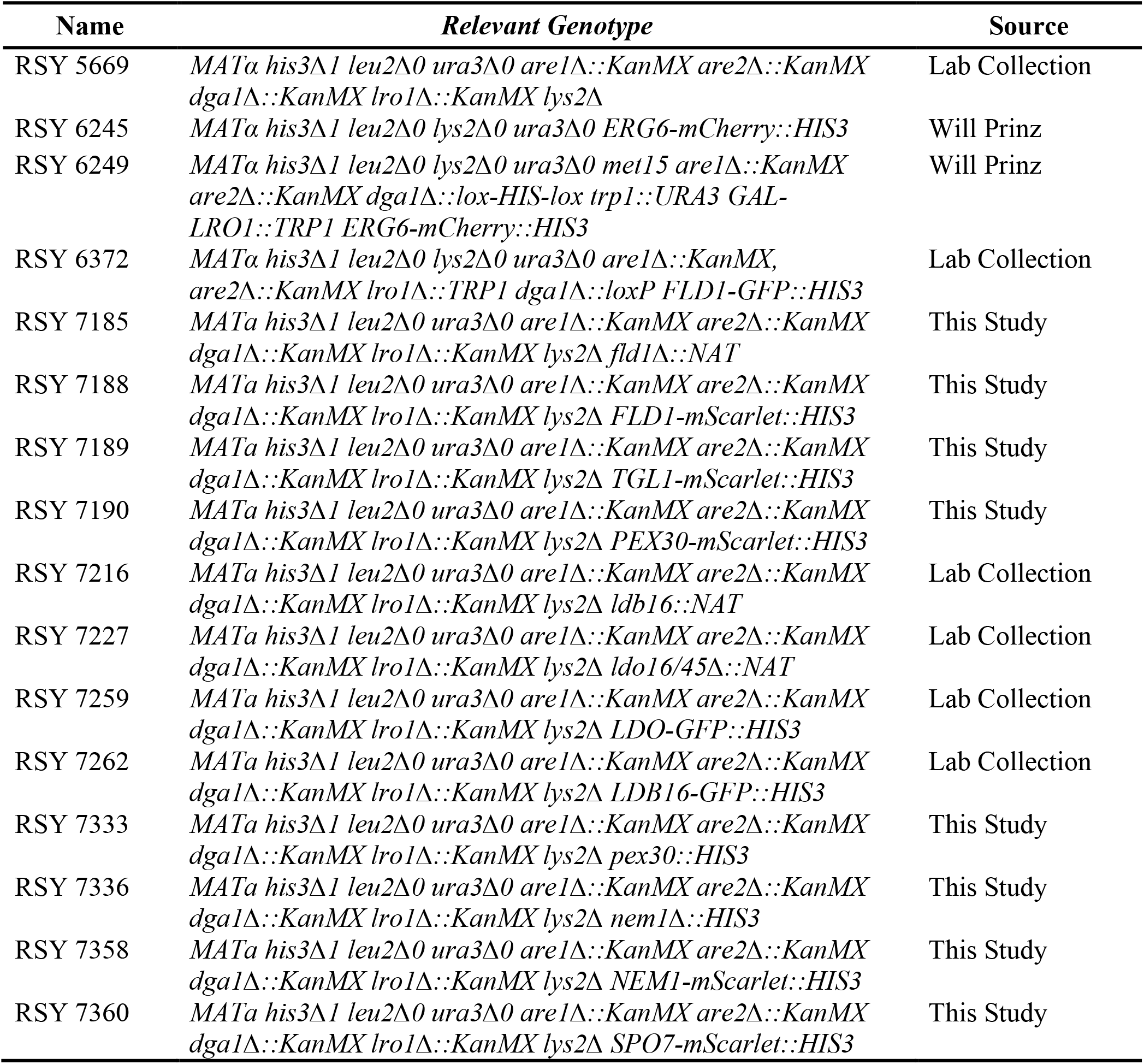
*Saccharomyces cerevisiae* strains used in this study

**Table S2.** Plasmids used in this study

| Plasmids | Source |
| --- | --- |
| pCM189-tetO<sub>7</sub>-PLIN3-GFP/URA3 | Khaddaj et al., 2022 |
| pCM189-tetO<sub>7</sub>-WBP1-GFP-PLIN3/ URA3 | Khaddaj et al., 2022 |
| pCM189-tetO<sub>7</sub>-SEC61-GFP-PLIN3/ URA3 | Khaddaj et al., 2022 |
| pCM189-tetO<sub>7</sub>-WBP1-GFP-PLIN3/ HIS3 | This Study |
| pCM189-tetO<sub>7</sub>-SEC61-GFP-PLIN3/ HIS3 | This Study |
| pCM189-tetO<sub>7</sub>-WBP1-GFP/ URA3 | Khaddaj et al., 2022 |
| pCM189-tetO<sub>7</sub>-SEC61-GFP/ URA3 | Khaddaj et al., 2022 |
| pCM189-tetO<sub>7</sub>-WBP1-mScarlet-PLIN3/ URA3 | Khaddaj et al., 2022 |
| pCM189-tetO<sub>7</sub>-SEC61-mScarlet-PLIN3/ URA3 | Khaddaj et al., 2022 |
| pCM189-tetO<sub>7</sub>-WBP1-mScarlet-PLIN3/ LEU2 | This Study |
| pCM189-tetO<sub>7</sub>-SEC61-mScarlet-PLIN3/ LEU2 | This Study |
| pCM189-tetO<sub>7</sub>-WBP1-mScarlet/ URA3 | Khaddaj et al., 2022 |
| pCM189-tetO<sub>7</sub>-SEC61-mScarlet/ URA3 | Khaddaj et al., 2022 |
| pRS415-ADH-mCherry-HDEL/ LEU2 | Lab Collection |
| pCM189-tetO<sub>7</sub>-WBP1-GFP-PLIN3<sup>V158D</sup>/ URA3 | This Study |
| pCM189-tetO<sub>7</sub>-SEC61-GFP-PLIN3<sup>V158D</sup>/ URA3 | This Study |
| pCM189-tetO<sub>7</sub>-WBP1-mScarlet-PLIN1/ URA3 | This Study |
| pCM189-tetO<sub>7</sub>-SEC61-mScarlet-PLIN1/ URA3 | This Study |
| pCM189-tetO<sub>7</sub>-WBP1-mScarlet-PAT/ URA3 | This Study |
| pCM189-tetO<sub>7</sub>-WBP1-mScarlet-11-mer/ URA3 | This Study |
| pCM189-tetO<sub>7</sub>-WBP1-mScarlet-4-helix bundle/ URA3 | This Study |
| pCM189-tetO<sub>7</sub>-WBP1-mScarlet-PAT+11-mer/ URA3 | This Study |
| pCM189-tetO<sub>7</sub>-WBP1-mScarlet-11-mer+4-helix bundle/<br>URA3 | This Study |
| pCM189-tetO<sub>7</sub>-WBP1-mScarlet-PAT+4-helix bundle/<br>URA3 | This Study |
| pCM189-tetO<sub>7</sub>-SEC61-mScarlet-PAT/ URA3 | This Study |
| pCM189-tetO<sub>7</sub>-SEC61-mScarlet-11-mer/ URA3 | This Study |
| pCM189-tetO<sub>7</sub>-SEC61-mScarlet-4-helix bundle/ URA3 | This Study |
| pCM189-tetO<sub>7</sub>-SEC61-mScarlet-PAT+11-mer/ URA3 | This Study |
| pCM189-tetO<sub>7</sub>-SEC61-mScarlet-11-mer+4-helix bundle/<br>URA3 | This Study |
| pCM189-tetO<sub>7</sub>-SEC61-mScarlet-PAT+4-helix bundle/<br>URA3 | This Study |
| pYEplac181-ADH-ER-DAG-Sensor/ LEU2 | Will Prinz |
| pRS416-ADH-TGL4-GFP/ URA3 | Lab Collection |
| PRS415-ADH-CDS1/ LEU2 | Lab Collection |
| PRS415-ADH-DGK1/ LEU2 | Lab Collection |
| pRS415-GAL-AYR1-GFP/ LEU2 | Lab Collection |
| pGREG503-PAH1-7A/ HIS3 | Lab Collection |

